# Metabolic Perturbation Exacerbates Sinoatrial Node Dysfunction in Heart Failure

**DOI:** 10.64898/2026.02.06.704519

**Authors:** Lu Ren, Yu Liu, Daphne A. Diloretto, Mengjie Xie, Yuting Pan, Yang Zheng, Arianne Caudal, Wenshu Zeng, Ryan L. Woltz, Danielle H Shin, Yuanyuan Dai, Rosmery Rodriguez, Richard Q. Ngo, Guy A. Perkins, Gabriela Grigorean, Ebenezer N. Yamoah, Joseph C. Wu, Hao Zhang, Nipavan Chiamvimonvat, Phung N. Thai

## Abstract

Heart failure (HF) affects approximately 6.2 million people in the United States, with a 5-year mortality exceeding 50%. Bradyarrhythmia, a known complication in HF due to sinoatrial node (SAN) dysfunction (SAND), increases morbidity and mortality of HF patients. Insights into the mechanistic underpinnings of SAND in HF could therefore uncover vital therapeutic targets to improve clinical outcomes. The SAN is endowed with a dense mitochondrial network crucial for sustaining their pacemaking function on a beat-to-beat basis. We have previously demonstrated significant disruptions in the mitochondrial-sarcoplasmic reticulum connectomics, resulting in abnormal mitochondrial Ca^2+^ handling and impaired mitochondrial function in HF. Here, we hypothesize that the metabolic perturbation is one of the critical mechanisms underlying SAND. To this end, we took advantage of a multi-omics approach combined with ultra-resolution imaging and functional analyses to decipher the metabolic shift that transpires in the HF SAN. Our findings revealed significant metabolic remodeling within the SAN mitochondria in HF, associated with altered fatty acid β-oxidation, ketone bodies, and carbohydrate catabolism, together with heightened mitochondrial cristae remodeling. Notably, metabolomics analyses identified pronounced increase of glucosylceramides and ceramides, motivating functional studies in human iPSC-derived SAN cells (iSANCs). Mixed ceramide exposure reduced stimulated oxygen consumption and the ATP-linked respiratory endpoint, increased extracellular acidification, and dose-dependently diminished spontaneous beating frequency. Collectively, our findings support the notion that ceramides are active mediators of mitochondrial and metabolic dysfunction in the SAN. Overall, the study provides evidence that ceramides may be a potential therapeutic target for mitigating SAND in HF.

## INTRODUCTION

Heart Failure (HF) is a progressive disease that impacts more than 23 million people worldwide.^1^ Despite recent advancements in therapeutic options for HF, the 5-year mortality remains high at 45-60%.^1,2^ A known complication of HF is bradyarrhythmia due to sinoatrial node (SAN) dysfunction (SAND), which exacerbates the already debilitating clinical syndrome of HF and heightens the morbidity and mortality among HF patients. Indeed, approximately 20-50% of sudden cardiac deaths in HF result from bradyarrhythmias, secondary to SAND.^3^ Elucidating the molecular and cellular mechanisms could therefore provide therapeutic targets for drug development and improve clinical outcomes.

Significant SAN remodeling occurs in HF, driven partially by elevated hemodynamic stress,^4^ increased neurohormonal activation,^5^ and alterations in the extracellular environment.^6^ In HF patients, these maladaptive alterations translate into prolonged intrinsic sinus cycle length, extended SAN conduction time, and prolonged corrected sinus node recovery time, which ultimately impair intrinsic heart rate.^7^ Despite attempts by neurohormonal modulations to curtail the depressed SAN function, the extensive remodeling induces chronotropic incompetence, which impedes the ability of the dominant pacemaker to respond sufficiently.^8^ Indeed, the impairment in intrinsic SAN function is manifested in multiple animal models of HF,^8–12^ suggesting that SAND frequently occurs concomitantly with the development of HF.

Despite lacking robust contractile properties, the SAN is endowed with a dense network of mitochondria to support its pacemaking function on a beat-to-beat basis.^13^ As the Ca^2+^ clock and membrane voltage clock rely heavily on the reestablishment of ionic gradients with every heartbeat, it is no surprise that the healthy SAN requires a constant and substantial supply of energy to meet its metabolic needs. *However, a knowledge gap remains regarding the dominant energy source in the SAN under normal physiological conditions and whether a metabolic shift in energy utilization in the SAN in HF may compromise SAN function, as commonly observed in HF patients.* Indeed, in contractile cardiomyocytes, the regression of energy utilization to the fetal-like state is detrimental to the energy-starved heart.^14–16^ It is unknown whether energetic remodeling occurs in the SAN, and importantly, its impact on the SAN’s function.

We have previously demonstrated that direct communication between mitochondria and the sarcoplasmic reticulum (SR) is crucial in ensuring the proper coupling of energy demand and supply. Specifically, disruptions in the mitochondria-SR connectomics result in abnormal mitochondrial Ca^2+^ handling and impaired mitochondrial function in the SAN in a preclinical model of HF.^17^ The objective of the current study is to investigate the mechanistic underpinnings of mitochondrial remodeling and the consequences of mitochondrial dysfunction in SAND in HF. We hypothesize that metabolic abnormalities occur in the SAN due to the accumulation of critically impaired mitochondria, which promote SAN dysfunction in HF. To this end, we leveraged an integrative multi-omics approach to investigate changes in the transcriptome, proteome, and metabolome to decipher the metabolic disturbances that occur in the SAN in a preclinical model of HF. We utilized human-induced pluripotent stem cell-derived SAN cells (iSANCs) to test whether mixed ceramide exposure alters cellular respiration and spontaneous beating frequency.

To induce murine HF with reduced EF (HFrEF), mice were subjected to 8 weeks of pressure overload via transverse aortic constriction (TAC). Integrated multi-omics analyses on SAN tissues were performed and revealed altered abundance of protein and metabolites associated with fatty acid β-oxidation (FAO), carbohydrate metabolism, ketone metabolism, and redox regulation. In addition to the significant remodeling of the microenvironment and oxidative stress, we found significant elevations in glucosylceramides and ceramides. These observations motivated a second stage of experiments testing whether mixed ceramide (MC) exposure affects human iSANC respiration and automaticity. We found that pretreatment of iSANCs with MCs progressively impaired FAO while enhancing anaerobic respiration. Further, iSANCs pre-treated with MCs exhibited profound bradycardia in a dose-dependent manner. This is the first report of integrative multi-omics analyses to elucidate the mechanistic underpinnings of SAND in a preclinical model of HF. The findings provide novel insights into a potential mechanism of metabolic perturbation and have important clinical implications in the rational design of therapeutic development for SAND in HF.

## RESULTS

### HF mice exhibited sinoatrial node dysfunction (SAND)

We induced HF by pressure overload via transverse aortic constriction (TAC), as previously described.^17^ Male and female mice were randomly assigned to sham or TAC operation. Due to the small size of the SAN (<10 mg of tissue), we pooled *five SAN tissues per sample* for multi-omic analyses. We performed proteomics, metabolomics, single nuclei-RNA sequencing (snRNA-seq), EM tomography and functional metabolic and electrophysiological assessment to decipher the mechanism of SAND in HF (**Fig. 1A**). HF mice exhibited a significant rightward shift in R-R interval, as shown in the representative heart rate variability (HRV) and histogram plots (**Fig. 1B**). This bradycardic shift in HF was also apparent when the circadian rhythms was accounted for. Of note, sex-stratified analyses of HRV revealed that both male and female mice exhibited significant bradycardia following TAC, with no significant sex differences in RMSSD or SDNN (**SFig. 1**). Representative action potential recordings from isolated SANCs showed longer spontaneous cycle lengths after TAC, consistent with altered intrinsic pacemaker activity (**Fig. 1C**). These recordings directly assess intrinsic SANC automaticity but do not measure sinoatrial conduction time or sinus node recovery time. Importantly, we used conscious echocardiography to validate the HF model before further experimentation, evidenced by significant cardiac dilatation and systolic dysfunction, as shown in the short- and long-axis images (**Fig. 1D**) and M-mode traces (**Fig. 1E**), respectively. TAC mice exhibited significant bradycardia (**Fig. 1F**), left ventricular (LV) hypertrophy (**Figs. 1G-H**), dilatation (**Figs. 1I-J**), and systolic impairment (**Fig. 1K**). Additional cardiac structural and functional parameters are depicted in **SFig. 2**. Taken together, our preclinical model of HF demonstrates significant bradycardia, recapitulating the arrhythmic phenotype observed in HF patients.

**Figure 1.**
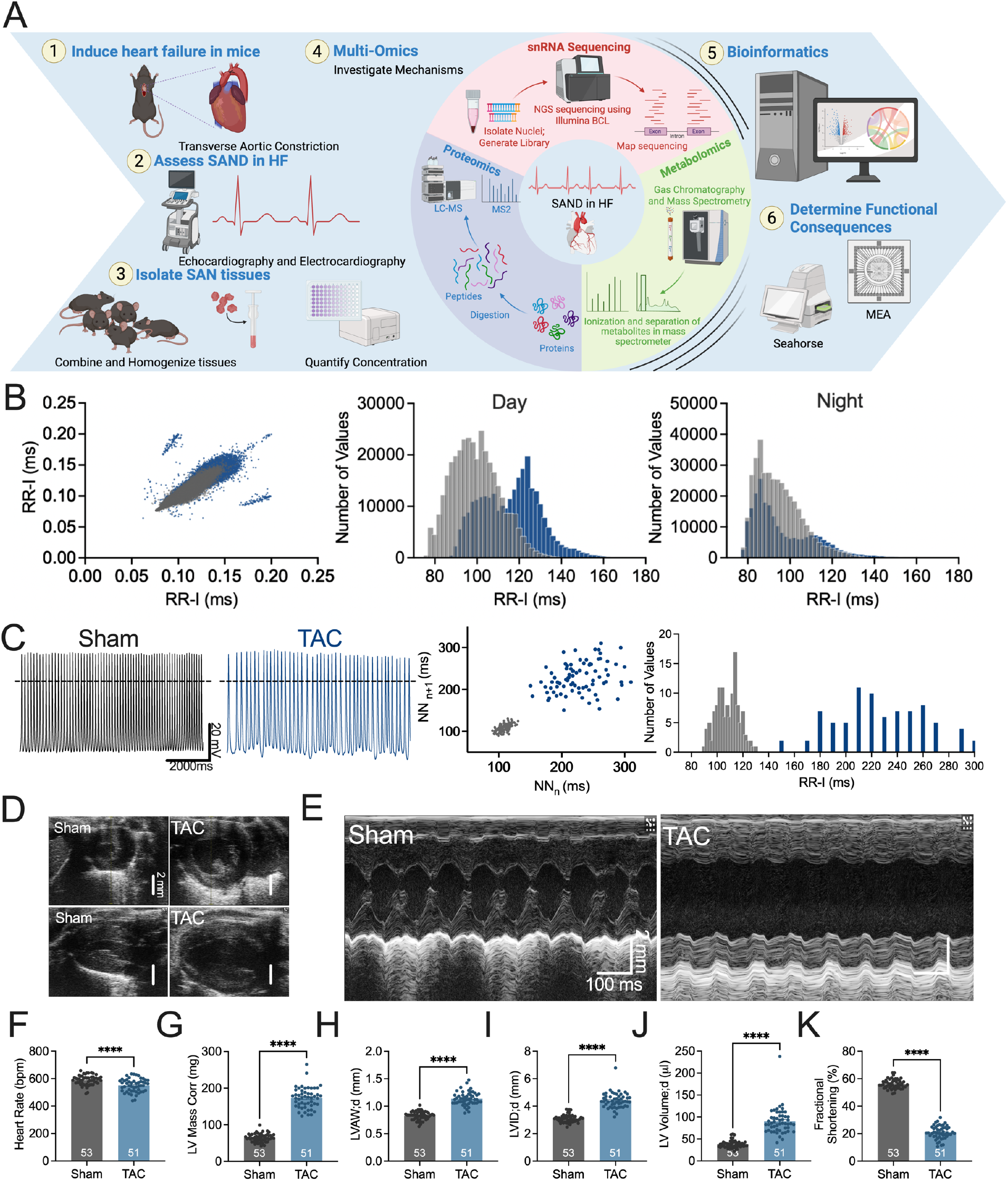
HF mice exhibited SAND. **A**) Graphical depiction of the overall design of the project. **B**) Representative Poincare and histogram plots of a sham and TAC mouse in the day and night time. **C**) Representative action potential recordings, Poincare, and histogram plots of both groups. **D**) Representative images of the short and long axes of the heart, and the **E**) M-mode images of both groups. Summary data of **F**) heart rate, **G**) left ventricular (LV) mass corrected, **H**) LV anterior wall end-diastolic thickness, **I**) LV internal diameter at diastole, **J**) LV end-diastolic volume, and **K**) fractional shortening. n=number of mice within bar graphs. n=number of mice, stated within bar graphs. Data expressed as mean±SEM. ****p<0.0001.

### Proteomic enrichment analysis revealed metabolic perturbations and elevated oxidative stress in the SAN of HF mice

We performed a comprehensive proteomic analysis in TAC and sham-operated mice to characterize the metabolic disturbances in the SAN in HF. The two groups showed distinct clustering (**SFig. 2**). Pathway enrichment analysis identified significant enrichment of several pathways related to metabolism, including general, protein, lipid, carbohydrate, and nucleotide metabolism, as depicted in the bubble plot (**Fig. 2A**). Rank analysis further revealed the importance of metabolic proteins within the SAN of both groups, highlighting Atp5f1a, a subunit of mitochondrial ATP synthase, as one of the top proteins in both groups (**Fig. 2B**). Moreover, the relative rank of the top 30 metabolic proteins considerably shifted in HF relative to control. The heatmap illustrates significant metabolic reprogramming, depicting the top 5 upregulated and 5 downregulated proteins in lipid, pyruvate, carbohydrate, glucose, vitamin, and protein metabolism, as well as proteins involved in the tricarboxylic acid (TCA) cycle and electron transport chain (ETC) (**Fig. 2C**). Significant changes across all metabolic pathways are provided in **STable 1**.

**Figure 2.**
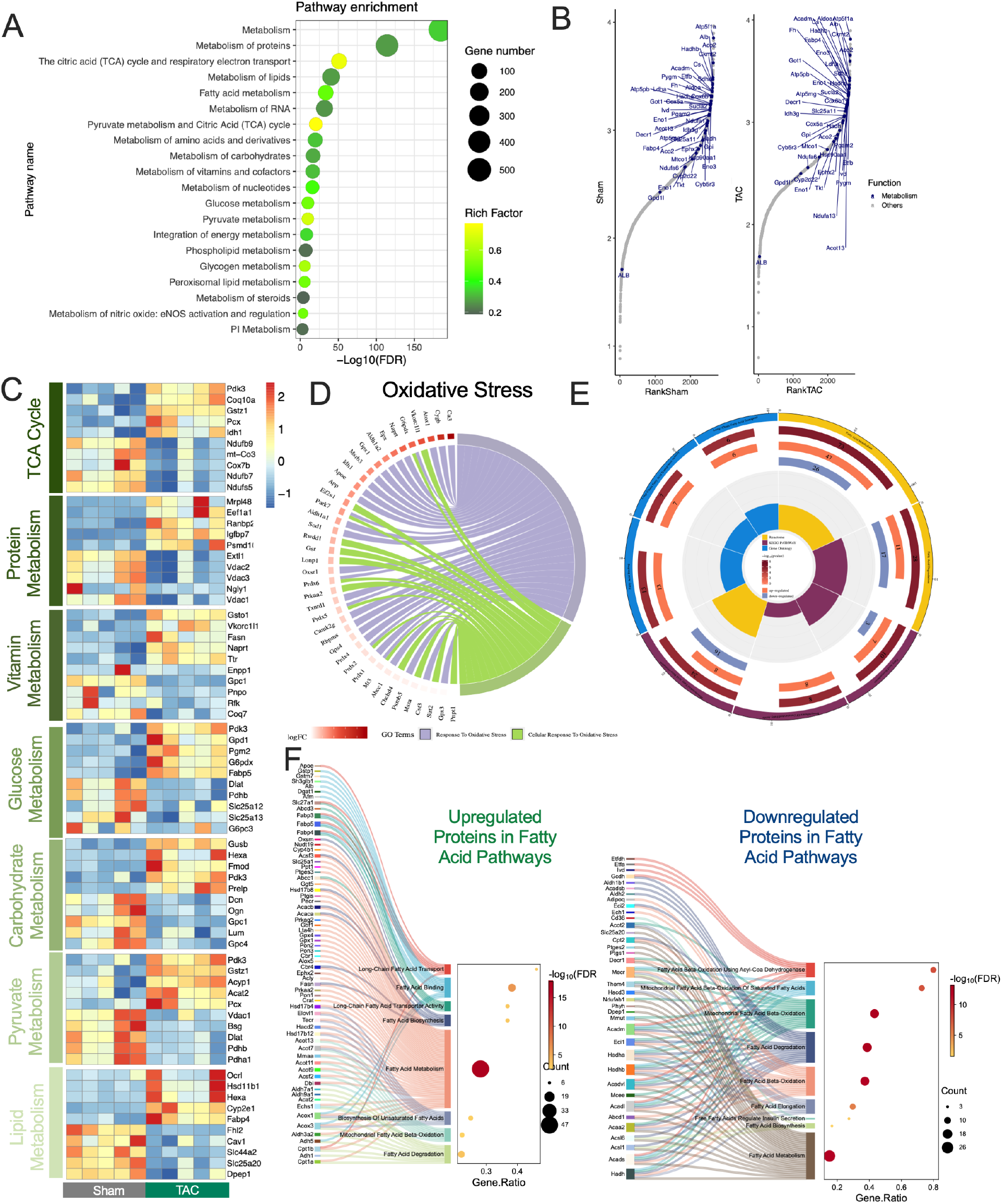
Proteomic enrichment analysis revealed metabolic perturbations in the SAN of HF mice. **A**) Bubble plot shows the top changes in the metabolic pathways enriched in the proteomics data. **B**) Abundance ranks of proteins within each group. Labeled proteins are involved in metabolism. **C**) Heat map shows the differential expression of proteins in the TCA cycle, as well as protein, vitamin, glucose, carbohydrate, pyruvate, and lipid metabolism in the SAN of sham and TAC mice. **D**) Chord plot shows the upregulated proteins in the oxidative stress pathway. **E**) Plot demonstrates the number of proteins upregulated and downregulated in fatty acid pathways. **F**) Graph shows the upregulated and downregulated proteins involved in multiple fatty acid pathways. n=5 pooled samples for each group; each pooled sample contains 4-5 SAN tissues. Data expressed as mean±SEM. *p<0.05, **p<0.01, ***p<0.001.

In the HF SAN, pyruvate dehydrogenase complex (PDC) subunits Pdha1, Pdhb, and Dlat were decreased. Among PDC regulatory kinases, Pdk1, Pdk2 and the cardiac-predominant Pdk4 were decreased, while Pdk3 was increased. Mitochondrial pyruvate carrier Mpc2 was also elevated.

In the oxidative pentose phosphate pathway (PPP), G6pdx, Pgls, and Pgd were all upregulated. Hexosamine and UDP-sugar pathway enzymes Uap1, Uap1l1, and Ugdh were increased, while O-GlcNAc transferase Ogt was decreased. Glycogen metabolism enzymes Ppp1r3a, Agl, Pygm, and Gys1 were uniformly decreased.

Alongside these metabolic changes, redox-related proteins showed compartment-specific remodeling in HF SAN. Mitochondrial antioxidant enzymes were decreased, including Sod2, Nadk2, Msrb2, Me3, Txn2, Prdx3, and Txnrd2. In contrast, cytosolic antioxidant enzymes Gsto1, Gpx1, and Ftl1 were increased. Chord plot analysis further identified upregulation of antioxidant enzyme families in HF SAN, including glutathione peroxidases (Gpx1, Gpx3, Gpx4), peroxiredoxins (Prdx1, Prdx5), thioredoxin reductase Txnrd1, superoxide dismutase Sod1, and catalase. Coenzyme Q (CoQ) biosynthesis enzymes Coq5, Coq7, and Coq9 were also decreased, while the CoQ regulatory kinase Coq8a was increased (**Stable 1**).

Complementing these redox alterations, lipid metabolism was broadly dysregulated in HF SAN (**Fig. 2E**). Fatty acid transporter Slc27a1, fatty acid binding proteins Fabp3, Fabp4, and Fabp5, and the acylcarnitine transferases Cpt1a and Cpt1b were all upregulated (**Fig. 2F**). In contrast, β-oxidation enzymes Acads, Acadm, Acadl, and Acadvl were decreased, along with downstream enzymes Hadh and Acaa2, and acyl-CoA synthetases Acsl1 and Acsl6. Carnitine-acylcarnitine translocase Slc25a20 and carnitine palmitoyltransferase 2 Cpt2 were also markedly decreased, consistent with altered mitochondrial fatty acid handling. Caveolins Cav1 and Cav3, structural proteins of sarcolemmal lipid rafts and caveolae, were among the most substantially decreased proteins in HF SAN. Overall, these data reveal broad metabolic remodeling in the HF SAN, characterized by altered abundance of proteins associated with fatty acid β-oxidation, carbohydrate metabolism, redox regulation, and ketone metabolism.

### The metabolomic profile demonstrated a significant upregulation of lipid metabolites

To investigate the metabolite-level consequences of these enzymatic shifts, we performed untargeted metabolomic analysis encompassing primary metabolism, biogenic amines, and lipidomics using pooled samples referenced against internal standards (**STable 2**). Principal component analysis demonstrated clear separation of sham and TAC SAN groups following normalization (**SFigs. 3–4**), and pathway enrichment identified purine metabolism, pantothenate and CoA biosynthesis, and acyl carnitine biosynthesis as the top enriched categories, respectively (**Fig. 3A**). STRING network and chord plot analyses revealed interrelationships among the altered metabolites and pathways (**Figs. 3B-C**).

**Figure 3.**
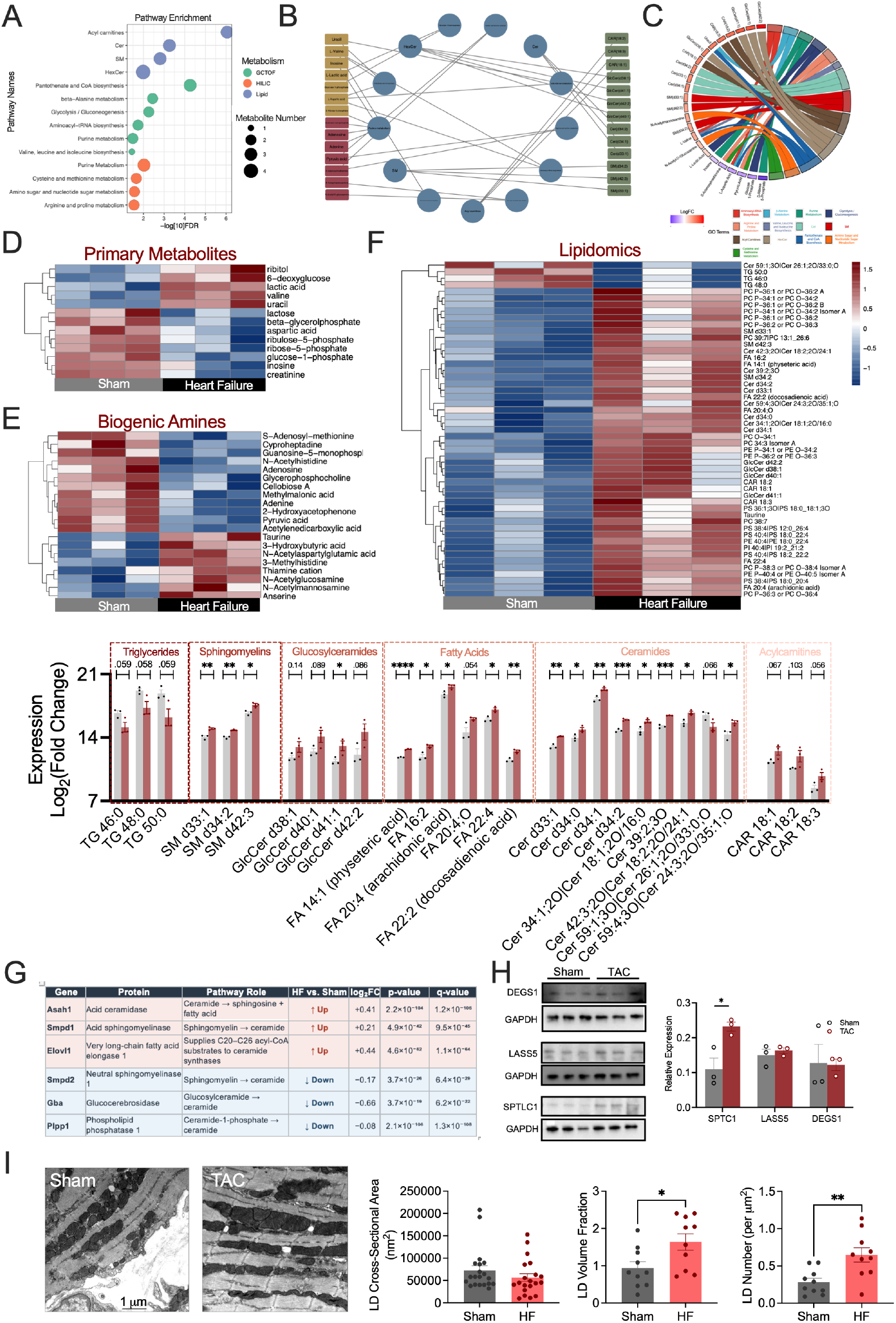
Metabolomic profile demonstrated significant upregulation of lipids. **A**) Bubble plot of the pathways enriched in the primary metabolites, biogenic amines, and lipidomics. **B**) STRING network analysis shows the interactions between the various metabolites that were altered. **C**) Chord plot shows the differential expression of metabolites and the pathways they are involved in. **D**) Heat maps show the differential expression of primary metabolites, **E**) biogenic amines, and **F**) lipids in sham and TAC SAN. Summary data of the lipids that were differentially expressed. **G**) Table showing ceramide proteins detected by proteomics and changed in HF. **H**) Representative western blot images and quantification of DEGS1, LASS5, and SPTLC1 protein expression in sham and TAC SAN tissue. **I**) Representative transmission electron microscopy (TEM) images of SAN tissue from Sham and TAC mice demonstrating lipid droplets (LD). Quantification of LD cross-sectional area (nm²), LD volume fraction (%), and LD number per µm² reveals a significant increase in LD volume fraction and LD number in TAC SAN compared to sham, with no significant difference in individual LD cross-sectional area. n=3 pooled samples for each metabolomics group, with each pooled sample contains 5 SAN tissues. Data expressed as mean±SEM. *p<0.05, **p<0.01, ***p<0.001, ****p<0.0001.

Glycolytic metabolites were broadly altered; 6-deoxyglucose and lactic acid were significantly elevated while glucose-1-phosphate and pyruvic acid were decreased (**Fig. 3D**). Valine, a branched-chain amino acid, was also elevated in the HF SAN, accompanied by an increase in methylmalonic acid. Pentose phosphate pathway (PPP) intermediates ribulose-5-phosphate and ribose-5-phosphate were downregulated despite increased PPP enzymes. 3-Hydroxybutyric acid was elevated alongside ketolytic enzymes, while adenosine and adenine were decreased.

Among biogenic amines, anserine, a histidine dipeptide with antioxidant and intracellular pH-buffering properties, was significantly elevated in HF SAN, corroborating the cytosolic antioxidant response observed in the proteome. Elevated 3-methylhistidine was consistent with altered protein turnover.

Beyond the shifts in carbohydrate and nucleotide metabolism, several lipid metabolites were significantly upregulated, highlighting lipid metabolism as a key altered pathway in HF SAN. Specifically, while triglycerides were decreased, there was an increase in sphingomyelins, glucosylceramides, fatty acids, ceramides, and acylcarnitines. A ranked summary of all significantly altered lipid species is provided in **STable 3**. Acylcarnitines (18:1, 18:2, and 18:3) were elevated alongside CPT1 (**Figs. 2, 3F**), while CPT2 and Slc25a20 were downregulated. Moreover, lipid metabolites such as phosphatidylcholine, phosphatidylserine, and phosphatidylethanolamine were significantly upregulated (**SFig. 6**). Among the elevated lipids, a remarkable number of ceramides exhibited a significant increase. Proteins detected in our proteomics data that are responsible for ceramide production and degradation are listed in **Fig. 3G**. Western blot analysis confirmed elevated SPTLC1, a core subunit of the rate-limiting enzyme in de novo ceramide synthesis serine palmitoyltransferase (SPT), in the HF SAN, while dihydroceramide desaturase 1 (DEGS1) and ceramide synthase 5 (LASS5) showed no significant change (**Fig. 3H**). TEM analysis showed greater lipid droplets in proximity to mitochondria without significant difference in individual droplet cross-sectional area in HF SAN (**Fig. 3I**). To characterize the molecular identity of these lipid droplets, we examined lipolysis enzymes, TG synthesis enzymes, and perilipin coat proteins by proteomics. While TG synthesis enzymes were unchanged, lipolytic enzymes HSL and MAGL were significantly upregulated, and the perilipin coat protein Plin1 (normally restricted to adipocytes) was significantly elevated in HF SAN (**STable 4**).

### Schematic of significant alterations in the proteome and metabolome in the SAN in HF

To visually examine the metabolic disruptions in the HF SAN, we mapped the significant alterations in the proteome and metabolome of key metabolic pathways, including glycolysis, the PPP, Krebs cycle, ketolysis, fatty acid *β*-oxidation, TCA cycle, ETC, as well as the transport mechanisms for glucose, fatty acids, and amino acids (**Fig. 4**). Upregulated proteins and metabolites were concentrated in the PPP, ketone handling and mitochondrial solute transport, while downregulated components were concentrated in fatty acid *β*-oxidation, TCA cycle, and ETC.

**Figure 4.**
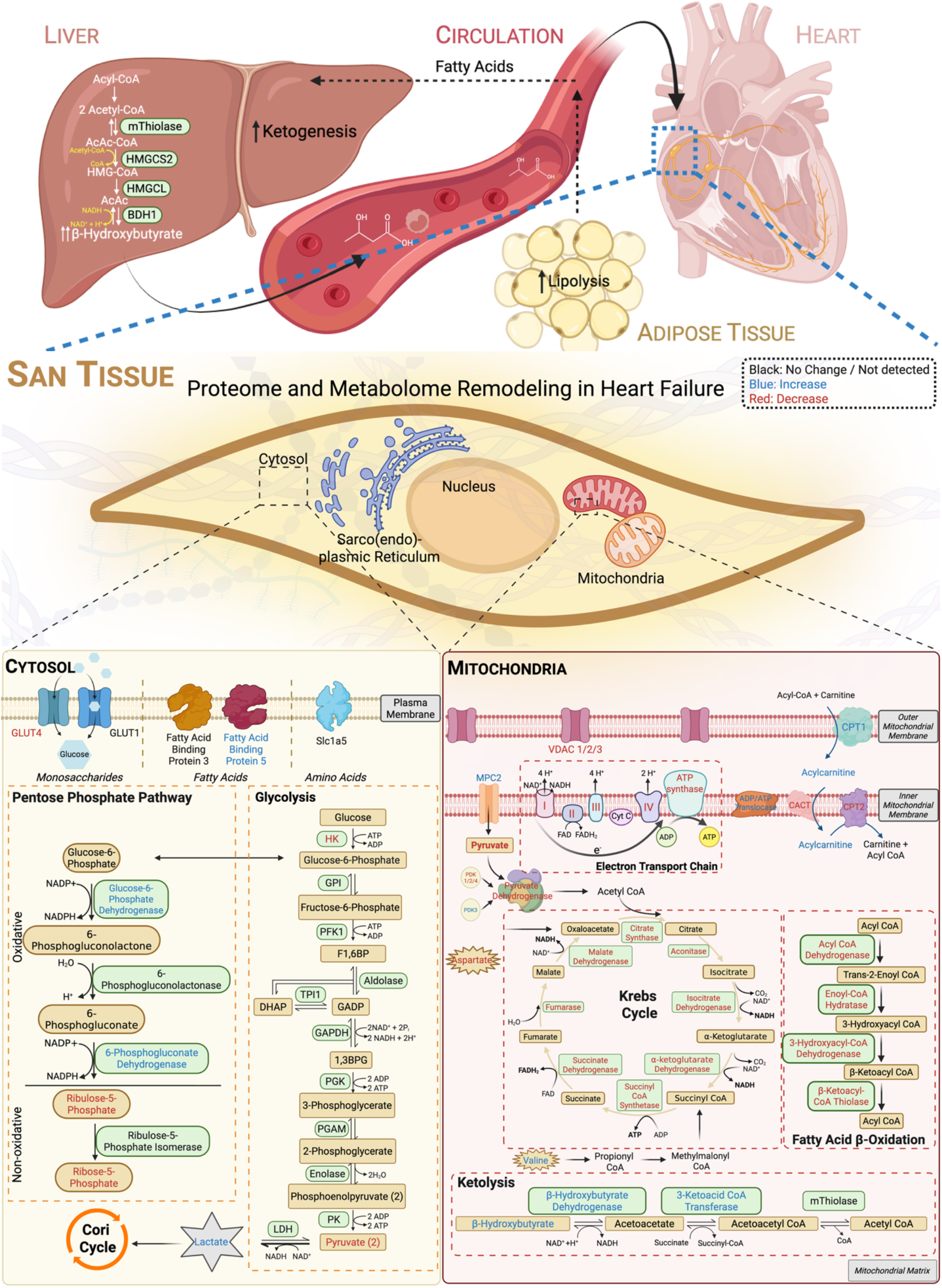
Schematic of significant alterations in the proteome and metabolome in the SAN in HF. Emphasis is placed on the changes seen in the proteins and metabolites involved in the major metabolic pathways in the SAN cell. Upregulated metabolites and proteins in HF SAN are represented in **blue**, while downregulated metabolites and proteins in HF SAN are represented in **red**. The pathways included in the graphical depiction of SAN metabolism include the pentose phosphate pathway, glycolysis, krebs cycle, fatty acid β-oxidation, ketolysis, and the transport of glucose, fatty acids, and amino acids.

To determine whether these metabolic alterations were specific to the SAN, we performed comparative multi-omic analyses using the SAN datasets generated in the current study, a previously published VCM proteomic dataset,^18^ and a newly generated VCM metabolomic dataset. Proteomics PCA (**Fig. 5A**) and metabolomic PCAs (**Fig. 5B**) showed distinct tissue- and condition-specific clustering. Cross-tissue proteomic comparisons identified specific and concordant protein changes (**Fig. 5C**). The SAN exhibited 505 upregulated and 305 downregulated proteins, whereas VCM showed 749 upregulated and 536 downregulated proteins (**Fig. 5D**). Metabolic module Z-scores (**Fig. 5E**), tissue divergence across 24 metabolic pathways (**Fig. 5F**), and key metabolic proteins (**Fig. 5G**) collectively revealed significant remodeling in both preparations. There was a markedly greater FAO-to-glycolysis shift in the HF SAN relative to VCM (**Fig. 5H**) and more severe median reduction in complex I and III subunit abundance in SAN, whereas the decrease in complex V was larger in VCM (**Fig. 5I**). Lipidomic comparison revealed SAN-specific accumulation of ceramides, free fatty acids, and polyunsaturated fatty acids not observed in VCM (**Figs. 5J-K**), alongside SAN-specific triglyceride saturation differences (**Fig. 5L**), cardiolipin remodeling (**Fig. 5M**), and primary metabolite alterations (**Fig. 5N**). Together, these findings demonstrate that the metabolic reprogramming in the HF SAN is distinct from that observed in VCM.

**Figure 5.**
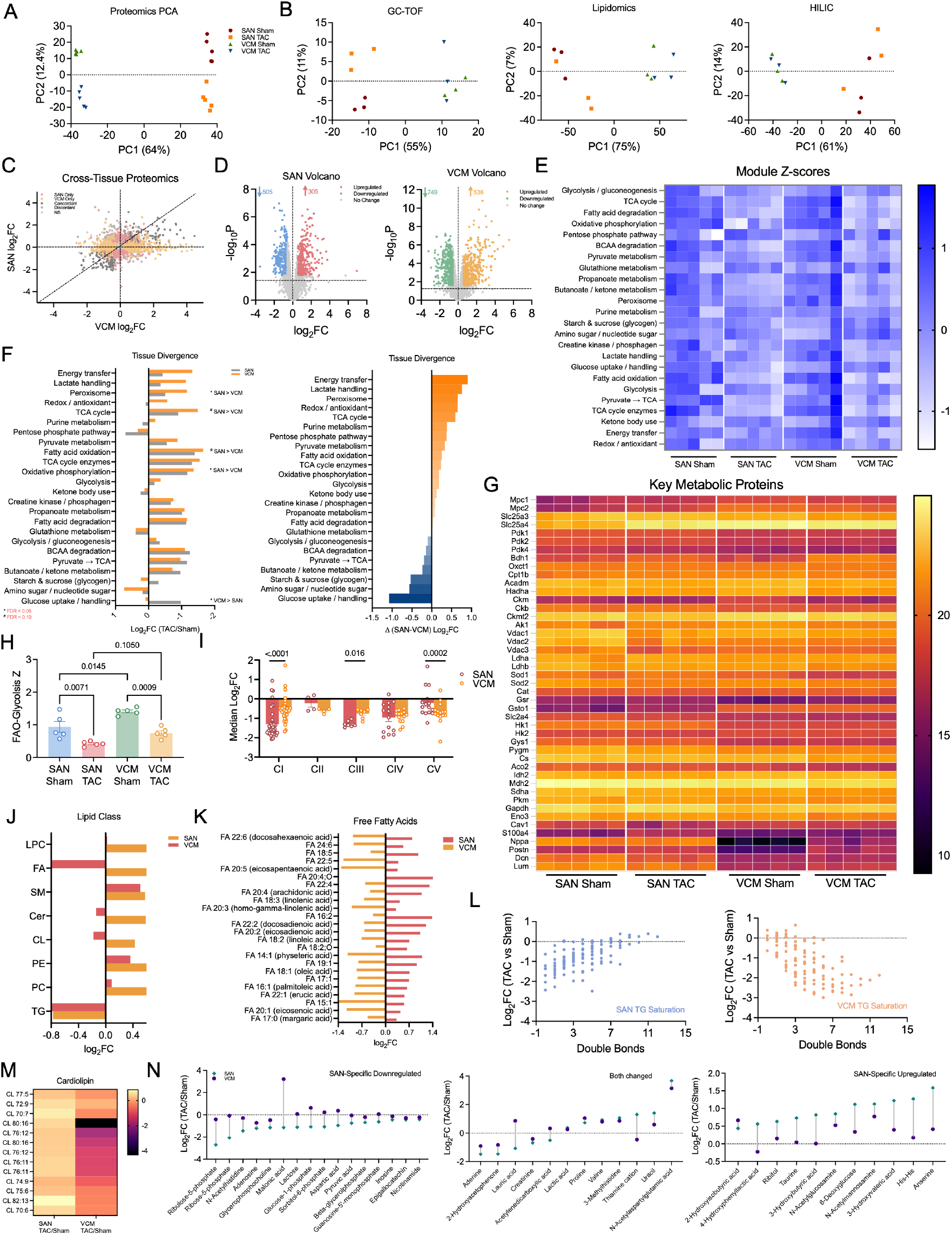
Comparative multi-omics analyses reveal distinct SAN-specific metabolic remodeling from ventricular cardiomyocyte adaptations in HF. **A)** Proteomic principal component analysis (PCA) SAN and ventricular cardiomyocyte (VCM) sham and TAC samples. PC1 separates tissues; PC2 separates conditions within each tissue. **B)** Metabolic PCAs for GC-TOF, HILIC, and lipidomics. **C**) Cross-tissue proteomics scatter plot comparing log₂FC (TAC/Sham) in SAN vs. VCM for all detected proteins. Points are classified as SAN-only, VCM-only, concordant, discordant, or non-significant. **D)** Volcano plots for SAN (505 upregulated, 305 downregulated) and VCM (749 upregulated, 536 downregulated). **E**) Metabolic module Z-score heatmap. **F**) Tissue divergence analysis showing log₂FC (TAC/Sham) for SAN and VCM across 24 metabolic modules (left), and the Δ(SAN−VCM) log₂FC (right). Modules with significant tissue divergence are annotated (* FDR < 0.05; # FDR < 0.10). **G**) Heatmap of 45 key metabolic proteins across all four groups. **H**) FAO-to-Glycolysis Z-score index across groups. **I**) Median log₂FC of OXPHOS subunits by complex (CI–CV) in SAN vs. VCM. **J)** Log₂FC for major lipid classes in SAN and VCM **K**) Log₂FC of individual free fatty acid species comparing SAN and VCM **L**) TG saturation analysis for SAN and VCM. **M**) Cardiolipin species heatmap (log₂FC, TAC/Sham) for SAN and VCM. **N**) Primary metabolite changes classified as SAN-specific downregulated, both changed, and SAN-specific upregulated, illustrating metabolomic features unique to the SAN in HF. Data expressed as mean±SEM. *p<0.05, **p<0.01, ***p<0.001, ****p<0.0001.

### snRNA-seq uncovered extensive remodeling of the external milieu

The SAN resides in a complex microenvironment composed of distinct cell types that work in concert to maintain the heart rhythm. To map cellular changes within the SAN in HF, we performed single-nucleus RNA sequencing. After sample processing and quality control, 55,249 nuclei were retained for gene expression analysis. Distinct clusters of adipocytes, B cells, cardiomyocytes (CMs), endothelial cells (ECs), fibroblasts (FBs), macrophages, neurons, and pericytes (PCs) were identified (**Fig. 6A**). Within the cardiomyocyte population, SANCs were distinguished from other cardiomyocytes by expression of Hcn1, Hcn2, Hcn4, Cntn2, Shox2, Cacna1h, and Tnt2. Condition-split UMAPs visualize the distribution of cell-type clusters separately in sham and TAC samples (**Fig. 6B**), with stacked bars showing the observed cell-type proportions in each pooled sample (**Fig. 6C**). The volcano plot shows the top 10 upregulated and downregulated genes in HF SAN (**Fig. 6D**), and violin plots depict the top 20 differentially expressed metabolic genes between the two groups (**SFig. 7**). Cell type identities were confirmed using established markers (**Fig. 6E**). Cell-type-resolved pathway analysis revealed distinct transcriptional responses across all populations (**Fig. 6F; SFig. 8**), including transcriptional signatures related to stress and hypertrophy in CMs, angiogenesis and extracellular matrix (ECM) remodeling in ECs, inflammation in immune cells, fibrogenic ECM remodeling in FBs, lipid metabolism in adipocytes, vascular remodeling in pericytes (PCs), and stress responses in neurons.

**Figure 6.**
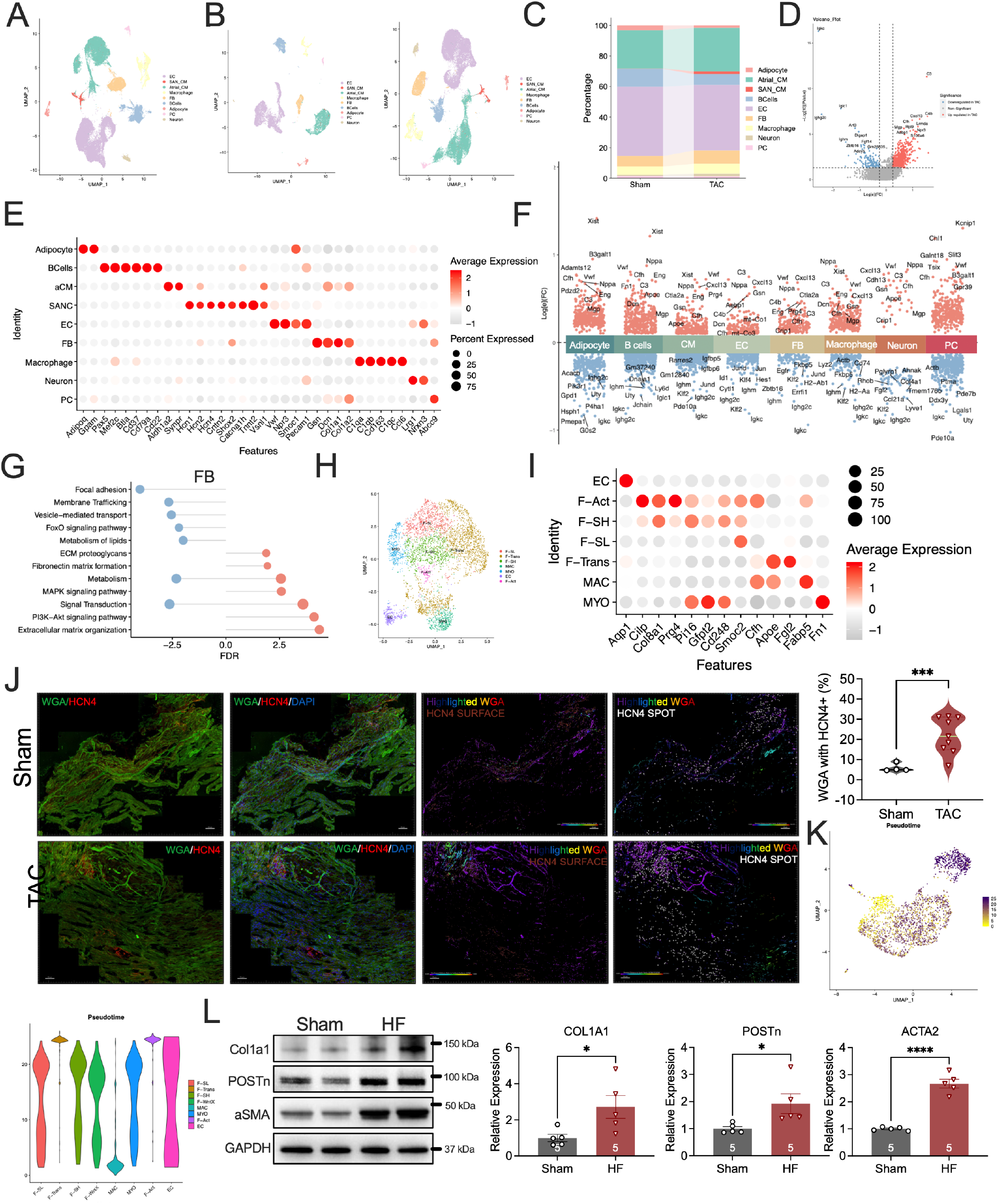
snRNA-seq uncovered extensive, adverse remodeling of the external milieu. **A**) The uniform manifold approximation and projection (UMAP) of isolated nuclei of SAN tissues of sham- and TAC-operated mice. **B**) Condition-split UMAPs displaying the distribution of cell type clusters in sham (left) and TAC (right) mice separately. **C**) Stacked bar chart depicting the proportional composition of each cell type, including adipocyte, B cells, cardiomyocyte (CM), endothelial cell (EC), fibroblast (FB), macrophage, neuron, and pericyte (PC) in the pooled sham and TAC samples. **D**) Volcano plot of the differentially expressed genes (DEGs) between the two different groups. **E**) Dot plot shows the markers used to distinguish the cell type. **F**) Volcano plot shows the DEGs of the different cell types. **G**) Graphs show the enriched pathways of FBs. **H**) UMAP of the different fibroblast population. **I**) Dot plot shows the marker genes used to identify distinct fibroblast populations, including activated fibroblasts (F-Act), fibroblasts in high (F-SH) and low (F-SL) states, transitioning fibroblasts (F-Trans), and myofibroblasts (MYO). **J**) Representative immunofluorescence images of SAN tissue from Sham (top) and TAC (bottom) mice co-stained with wheat germ agglutinin (WGA, green), HCN4 (red), and DAPI (blue). Summary data of WGA surface and HCN4+ areas to the right. **K**) Pseudotime of the distinct populations. **L**) Representative western blot images of collagen type I alpha 1 (Col1a1), periostin (POSTn), a-smooth muscle actin (αSMA), and glyceraldehyde 3-phosphate dehydrogenase (GAPDH) and summary data for each are shown. Data expressed as mean±SEM. *p<0.05, ***p<0.001, ****p<0.0001.

Among these, FBs exhibited the most extensive transcriptional remodeling (**Fig. 6G**). Distinct subpopulations—FB-Sca1low (F-SL), FB-Sca1^high^ (F-SH), F-WntX, myofibroblasts (MYOs), and F-activated FBs (F-Act)—were resolved in the UMAP (**Fig. 6H**) using established markers (**Fig. 6I**). Consistent with FB activation, WGA co-staining showed increased WGA-positive area in HF SAN tissue (**Fig. 6J**). Pseudotime analysis revealed divergent developmental trajectories (**Fig. 6K**). Macrophages appeared at the lower end; F-SL, F-SH, F-WntX, MYOs, and ECs displayed broad intermediate distributions; and F-Act cells were skewed toward the higher end. Western blot confirmed elevation of fibrotic markers Col1a1, periostin (POSTN), and α-smooth muscle actin (αSMA) (**Fig. 6L**). Overall, snRNA-seq revealed extensive transcriptional remodeling across multiple cell types in the HF SAN microenvironment, with fibroblasts showing the most extensive remodeling.

### SAN mitochondria from HF mice showed impaired cristae structure and energy production

Given the evidence of impairment in energy mismatch (**Figs. 2**–**5**) and the central role of mitochondria in energy production, we investigated whether the metabolic dysregulation identified in our multi-omics analyses was linked to mitochondrial structural changes using transmission electron microscopy (TEM) and EM tomography. As depicted in EM tomography images at multiple angles (**Fig. 7A**) and quantitatively (**Fig. 7B**), the cristae density in the mitochondria of HF SANCs was significantly reduced compared to sham controls, with increased fragmentation and remodeling from the typical lamellar form to smaller, fragmented structures with tubular extensions. When SAN mitochondria of similar volume were examined, mitochondria from HF SANCs clustered around the lower cristae density (**Fig. 7C**).

**Figure 7.**
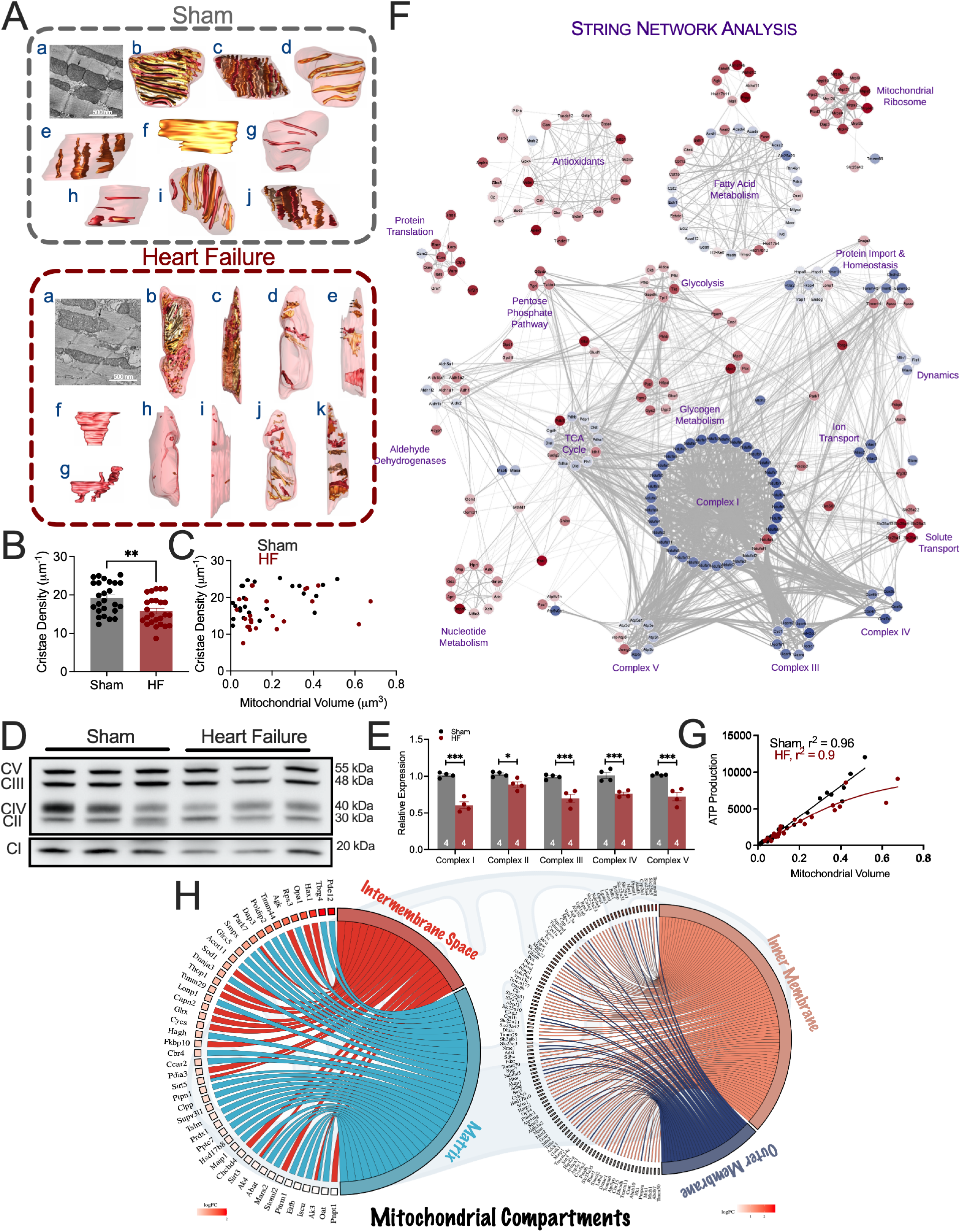
SAN mitochondria from HF mice showed impaired cristae structure and energy production. **A**) **Sham:** a) A representative slice through the middle of an EM tomography volume from sham SAN. The arrow points to the mitochondrion chosen for membrane segmentation. b) Top view of the surface-rendered mitochondrion after membrane segmentation. All 39 cristae are shown in various shades of brown, and the outer mitochondrial membrane (OMM) is shown in transparent maroon to better visualize the cristae. c) Side view of the mitochondrial volume. d) Top and e) side views of 5 representative cristae to show the typical lamellar nature of sham cristae and how they extend from side-to-side and top-to-bottom through the volume. f) A face of a typical lamellar sham crista that is in the orthogonal view relative to those shown in d and e. g) Top and h) side views of the 5 smallest cristae in this sham mitochondrion. i) Top and j) side views showing every third crista (13 out of 39). Displayed this way, the crista density is better compared between sham and TAC mitochondria because displaying the entire complement of cristae produces views in any orientation that obscure the perception of density due to cristae overlap. **Heart Failure:** a) A representative slice through the middle of an EM tomography volume from a SAN TAC mouse. The arrow points to the large mitochondrion chosen for membrane segmentation. b) Top view of the surface-rendered mitochondrion after membrane segmentation. All 52 cristae are shown (various shades of brown) and the OMM (translucent maroon). c) Side view of the mitochondrial volume. d) Top and e) side views of 5 representative cristae. Arrows point to the cristae that have been remodeled, showing a portion of their lamelli form becoming tubular extensions towards the mitochondrial periphery. f) A face of a lamellar TAC crista that is unlike the typical lamellar cristae of sham mitochondria. g) A face of a commonly seen TAC crista morphotype that shows remodeling to a smaller lamellar compartment and tubes extending above and below it. h) Top and i) side views of the 5 smallest cristae in this TAC mitochondrion. They are much smaller than the 5 smallest cristae in the sham mitochondrion. All 5 are small tubes suggesting not only cristae remodeling to the tubular form, but also cristae fragmentation. j) Top and k) side views showing every third crista (17 out of 52). Summary data showing **B**) cristae density and **C**) cristae density relative to mitochondrial volume. **D**) Representative western blot images of the five different complexes in the ETC and **E**) the associated summary data. **F**) STRING network of major metabolic pathways related to the mitochondria, based on MitoCarta. **G**) Summary data show modeled ATP production relative to mitochondrial volume in sham and HF groups. **H**) Chord plots show the differential expression of key proteins in the various compartments of the mitochondria, including the intermembrane space, matrix, inner membrane, and outer membrane. Data expressed as mean±SEM. *p<0.05, **p<0.01, ***p<0.001.

Protein expression levels of ETC complexes I-IV (CI-IV) were examined, and significant decreases were observed in all complexes (**Figs. 7D-E**). We performed STRING analysis on our proteomics data to investigate the specific changes that occurred (**Fig. 7F**). As depicted, key components of the energy production machinery, notably in the TCA cycle, ETC, and fatty acid oxidation, showed altered abundance. In particular, of the proteins detected in our proteomic dataset for CI, CII, CIII, CIV, and CV, 32/34, 1/2, 8/8, 5/5, and 6/8 were decreased, respectively, which corroborated with our western blot analyses of the diminished expression of the various complexes. In the TCA cycle, 9/12 of the proteins detected were declined, suggesting a depressed capacity to produce reduced equivalents for the ETC optimally. Conversely, pathways involved in mitochondrial ribosome, protein translation, solute transport, glycolysis, glycogen metabolism, and nucleotide metabolism were significantly upregulated. As previously determined (Fig. 2), antioxidant enzymes were upregulated, indicating an overall redox remodeling.

We further leveraged structural modeling of the mitochondria^19,20^ to estimate the rate of ATP production, based solely on structural parameters, regardless of the expression levels of enzymes involved in energy production (**Fig. 7G**). The modeled rate of ATP production was lower in the mid to large size HF SAN mitochondria (volume>0.2 μm^3^) relative to sham mitochondria, while it remained similar in the smallest mitochondria (volume<0.2 μm^3^).

In contrast, several critical proteins in different compartments of the mitochondria, including the intermembrane space, matrix, inner membrane, and outer membrane, were significantly upregulated in HF SAN (**Fig. 7H**). Notably, a few of the top upregulated proteins include ADP/ATP translocase 2 (Slc25a5), ADP/ATP translocase 4 (Slc15a31), and adenine nucleotide translocase type 1 (ANT1, Slc25a4), which serve to transport ATP to the cytosol. Overall, HF SAN mitochondria exhibited reduced cristae density, decreased ETC complex expression, lower modeled ATP production rates, and concurrent upregulation of mitochondrial ATP transport proteins.

### Mixed ceramides promoted a glycolytic shift, suppressed fatty acid β-oxidation, and impaired electrophysiological function in iSANCs

Since we observed an increase in ceramides alongside a metabolic shift, we hypothesized that ceramides may drive the changes in cellular respiration, given their established role in regulating metabolism. To directly test the hypothesis, we differentiated human iSANCs using well-established protocols and used this platform to test the effects of exogenous ceramides on iSANC respiration and automaticity (**Fig. 8A**). We first validated the pacemaker properties of iPSC-SAN cells relative to ventricular iPSC-derived cardiomyocytes (iPSC-VMs) using the patch-clamp technique in current-clamp configuration to ensure our model was reliable for mechanistic studies. iPSC-SAN cells exhibited a slower maximum upstroke velocity (dV/dt_max_) and a more depolarized maximum diastolic potential compared to iPSC-VMs (**Fig. 8B**). Moreover, iPSC-SANs showed reduced action potential duration (APD) ratios between 30-40 and 70-80% of repolarization (APD_30-40_/APD_70-80_), consistent with triangulation of AP morphology and their nodal status. At the molecular level, our quantitative PCR showed that iPSC-SANs exhibited higher expression of SAN markers including Shox2, T-box transcription factor (Tbx)18, Tbx5, and insulin enhancer protein (ISL)-1 relative to iPSC-VMS (**Fig. 8C**). Immunofluorescence imaging also demonstrated heightened protein abundance of Shox2, as well as greater hyperpolarization-activated cyclic nucleotide-gated channel 4 (HCN4) expression in iPSC-SAN cells (**Fig. 8D**).

**Figure 8.**
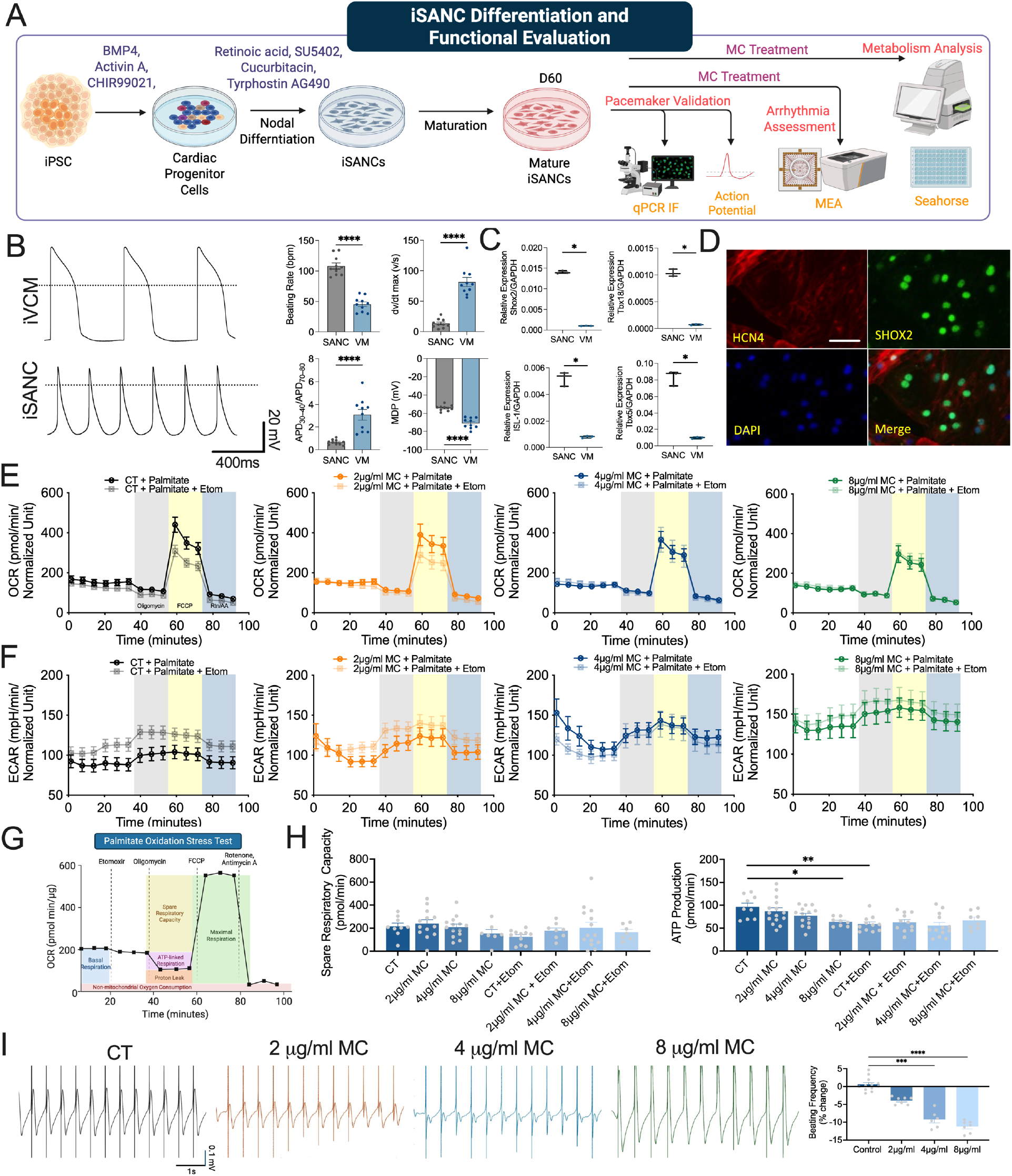
Mixed ceramides altered respiration and reduced spontaneous beating frequency in iSANCs. **A**) Protocol of iSANC differentiation and experimentation. **B**) Electrophysiological characterization of iPSC-SAN cells (iSANCs) compared to iPSC-derived ventricular cardiomyocytes (iPSC-VMs). Beating rate, dv/dt_max_, APD_30-40_/APD_70-80_, MDP (maximum diastolic potential) were assessed using patch clamp recordings. **p<0.01 by unpaired t-test. **C**) Gene expression analysis of key pacemaker markers (SHOX2, TBX18, ISL1, TBX5) in iPSC-SAN cells compared to iPSC-VMs. *p<0.05 by unpaired t-test. **D**) Immunofluorescence staining for the pacemaker markers Shox2 in green and hyperpolarization-activated cyclic nucleotide-gated channel 4 (HCN4) in red. DAPI in blue was used for nuclei staining. **E**) Summary data of oxygen consumption rate (OCR) in control, 2 μg/ml MC, 4 μg/ml MC, and 8 μg/ml MC treated iSANCs. **F**) Summary data of ECAR in control, 2μg/ml MC, 4 μg/ml MC, and 8μ g/ml MC treated iSANCs. **G**) Graphical depiction of the different parameters that were calculated form our data. **H**) Bar graphs show summary data of spare respiratory capacity and ATP production**. I**) Representative MEA recordings of the four different groups and summary data of the data. Data expressed as mean±SEM. *p<0.05, **p<0.01. ***p<0.001, and ****p<0.0001.

iSANCs were seeded onto 96-well plates and randomly pre-treated with either vehicle control (CT) or mixed ceramides (MCs) at increasing concentrations (2 µg/ml, 4 µg/ml, and 8 µg/ml) for 2 weeks. MCs were chosen based on the lipid-class increase in a broad range of ceramides in our metabolomics data. All wells contained palmitate as a substrate for fatty acid β-oxidation. Basal oxygen consumption rate (OCR) was comparable in all groups. As expected, OCR declined following oligomycin application in all conditions (**Fig. 8E**). Carbonyl cyanide 4-(trifluoromethoxy)phenylhydrazone (FCCP, a potent uncoupler of oxidative phosphorylation) application led to an OCR increase in all groups, with the control group exhibiting the most pronounced rise. In contrast, the 8 µg/ml MC group showed the least elevation. Furthermore, in the wells with ethyl-2-[6-(4-chlorophenoxy)hexyl]oxirane-2-carboxylate (ETOM), a CPT1 inhibitor, only the control group showed a noticeable decline in OCR after FCCP application. The application of rotenone/antimycin A reduced the OCR in all groups as expected. Moreover, our extracellular acidification rate (ECAR) data revealed that iSANCs treated with MCs dose-dependently xhibited increased extracellular acidification compared to control (**Fig. 8F**). Additionally, ETOM treatment elevated ECAR in control across all conditions but had no effect in MC-treated groups (**Fig. 8F**). We further calculated the spare respiratory capacity and ATP-linked respiratory endpoint (**Fig. 8G**). Although there was no difference in spare respiratory capacity, ATP production was significantly reduced in the 8 µg/ml MC group relative to the control, similar to the effects of ETOM on the control group (**Fig. 8H**).

To assess the functional consequences of MC exposure, we recorded baseline MEAs in iSANCs and then subjected them to either vehicle or various concentrations of MCs. As depicted in the representative images and summary data, beating frequency remained unchanged in the CT group (**Fig. 8I**). In contrast, treatment with MC inhibitors led to a dose-dependent decline in spontaneous beating frequency. Together, MC treatment dose-dependently decreased OCR and increased ECAR, reduced ATP production, and decreased spontaneous beating frequency.

## DISCUSSION

Sinoatrial node dysfunction (SAND) exacerbates the clinical outcome of HF patients, yet currently, no therapeutics exist to correct the impaired intrinsic function of the SAN in HF. This is primarily attributed to a substantial knowledge gap in our mechanistic understanding of SAND in HF. To bridge this gap and address the unmet clinical need, we applied a comprehensive approach combining multi-omics analyses, ultra-resolution imaging using EM tomography, and metabolic and functional assessment of iSANCs to test the hypothesis directly. Indeed, this study represents the first to identify the metabolic remodeling that may contribute to SAND in HF with reduced ejection fraction (HFrEF). Our findings highlight the central role of the mitochondria in SAN automaticity in both health and disease and provide evidence for the role of ceramides as a potential therapeutic target to mitigate metabolic disturbances and alleviate SAND.

### Significant alterations in the ultrastructure of mitochondria in HF SAN

As the primary energy-producing organelles, the proper functioning of mitochondria is crucial for sustaining SAN automaticity. We found extensive structural remodeling in the mitochondria from HF SAN, particularly at the inner mitochondrial membrane, where the ETC machinery resides (**Fig. 7**). The mitochondria from HF SAN exhibited a transition to more branched and condensed mitochondria^17^ with a concomitant decline in cristae density, potentially accounting for the 10-40% reduction in the complexes of the ETC. Despite the overall decrease in expression of these critical components of the energy-producing machinery, the ADP/ATP translocases were significantly upregulated, likely a compensatory response to maintain sufficient ATP levels in the cytosol. Consistent with the current findings, our previous study demonstrated that SANCs from HF mice exhibit diminished ATP production when perfused with glutamate/malate and succinate to stimulate CI and CII, respectively.^17^ These earlier findings corroborate the structural modeling data, which predict lower ATP production in HF SAN with accounted mitochondrial volume (**Fig. 7**).

### Metabolic shift in energy generation

Our current study not only identifies the possible metabolic preference of the murine SAN in health, but it has also demonstrated multifaceted metabolic perturbations that promote the uncoupling of energy demand and supply in the SAN in HF. Indeed, multi-omics analyses revealed significant disruption in the lipidomic profile in the HF SAN. Specifically, CPT1, the rate-limiting enzyme in fatty acid β-oxidation was significantly upregulated, but the downstream enzymes required to catalyze the subsequent steps within the mitochondrial matrix exhibited reduced expression. Moreover, the critical enzymes essential for the medium-, long-chain-, and very long-chain fatty acid β-oxidation, notably Acadm, Acadl, and Acadvl, were significantly reduced, potentially contributing to the accumulation of fatty acids in lipid droplets located near the mitochondria (**Fig. 3**). Maintaining lipid homeostasis is critical for normal cellular function. To mitigate the risk of lipotoxicity, cells sequester lipids into droplets that may appear near the mitochondria.^21^ Proteomic interrogation of canonical lipolytic enzymes (ATGL/Pnpla2, HSL/Lipe) and triglyceride synthesis enzymes (DGAT1/2, AGPAT1-3, GPAM) revealed no significant changes in either pathway, indicating that LD accumulation reflects substrate overflow from impaired mitochondrial oxidation rather than induction of discrete anabolic or catabolic programs.^15,22^ Consistent with a pathological rather than physiological LD phenotype, the perilipin coat protein Plin1, normally an adipocyte marker of stable lipid storage, was significantly upregulated in HF SAN, while Plin4 was reduced, a pattern atypical for cardiac tissue, supporting lipotoxic sequestration rather than the regulated, oxidation-coupled LD buffering described in physiological contexts.^23^ Despite the spatial localization of lipids, the mitochondria in the SANCs are unable to efficiently utilize fatty acids, likely due to the downregulation of critical enzymes involved in lipid metabolism.

### A significant shift toward ketone body utilization and carbohydrate metabolism in HF SAN

This impaired lipid utilization is compensated by a metabolic shift toward greater reliance on ketone bodies and carbohydrates, as evidenced by the upregulation of critical enzymes and metabolites involved in these pathways (**Fig. 4**). In HF, hepatic ketogenesis is significantly upregulated, leading to elevated circulating ketone bodies,^24,25^ which not only affect contractile cardiomyocytes, but, as shown in this study, also impact the SAN (**Fig. 8**). As mitochondrial function and fatty acid β-oxidation become impaired, ketolysis and glycolysis take on a dominant role in cellular energy production. However, because carbohydrate and ketone body metabolism yield less ATP per weight than fatty acid β-oxidation,^15^ the detrimental shift may be energetically insufficient to meet the high demands of the SAN. We also observed increased valine in the HF SAN, suggesting altered branched-chain amino acid metabolism. Valine-derived carbon can contribute to the TCA cycle through its canonical conversion to propionyl-CoA and subsequently succinyl-CoA, while isotope-tracing studies have also demonstrated contributions to the acetyl-CoA pool.^26^ Akin to contractile cardiomyocytes, our data suggest that the SAN preferentially utilizes fatty acids in the healthy state and switches to carbohydrate/ketone body metabolism in HF.^15^ Indeed, contractile cardiomyocytes undergo genetic reprogramming to a fetal-like stage in HF, with increased dependence on carbohydrate metabolism.^27^ Despite its directional similarity, the magnitude of the shift and the associated lipid profile changes differ substantially (**Fig. 5**). To our knowledge, this study provides the first evidence suggesting a metabolic shift in the SAN, illuminating the importance of the *energy starvation hypothesis*^14,15,28^ to pacemaking cardiomyocytes.

### Ceramides compromise mitochondrial function

In parallel with the metabolic perturbation, we observed substantial accumulation of glucosylceramides and ceramides in the HF SAN (**Fig. 3**). Ceramides are bioactive signaling molecules implicated in the pathogenesis of cardiac diseases, including diabetic cardiomyopathy, coronary heart disease, and HF, with their expression level correlating positively with disease severity.^29,30^ Pharmacological inhibition of ceramides, or genetic knockout of enzymes that catalyze the synthesis of ceramides, has been shown to be cardioprotective in multiple cardiovascular diseases.^31^ This suggests that elevated levels of ceramides may be detrimental to cellular survival. Indeed, increased expression of ceramides has been shown to induce oxidative stress,^40,41^ impair fatty acid oxidation,^32,33^ disrupt ETC activity,^34^ and lead to overall mitochondrial dysfunction.^35^

To elucidate the source of ceramide accumulation in the HF SAN, we examined key enzymes across the ceramide synthesis and degradation pathways by proteomics and Western blot (**Fig. 3**). Of the three enzymes examined by Western blot, SPTLC1, a core subunit of the rate-limiting enzyme in de novo ceramide biosynthesis serine palmitoyltransferase (SPT) was significantly upregulated in the HF SAN. As fatty acid β-oxidation fails, excess lipid substrates may be preferentially channeled into de novo sphingolipid synthesis via SPT, a phenomenon consistent with ceramide accumulation in lipotoxic states more broadly. Collectively, the proteomics data summarized in **Fig. 3** provide a comprehensive view of ceramide pathway remodeling in the HF SAN, with selective de novo biosynthesis upregulation emerging as the primary contributor to the ceramide burden.

As critical metabolic regulators, ceramides are known to antagonize insulin signaling by interfering with the activation of Akt/PKB, a crucial mediator of insulin’s metabolic effects.^36^ This interference ultimately reduces insulin-stimulated translocation of GLUT4, the primary glucose transporter in cardiac tissues, thereby diminishing insulin-dependent GLUT4 glucose uptake.^37^ Proteomics identified decrease of GLUT4 in the HF SAN; the change aligns with a prior study showing ceramide-mediateed suppression of insulin/Akt signaling that consequently impairs GLUT4 membrane translocation;^37^ GLUT1 upregulation trend likely represents a partial compensatory response to sustain basal, insulin-independent glucose uptake.

In the HF SAN, ceramide accumulation occurs alongside broad disruption of oxidative metabolism. Ceramides are established inhibitors of the mitochondrial respiratory chain,^38,39^ limiting electron transport capacity and promoting a compensatory shift toward glycolytic ATP production. Ceramide-mediated respiratory chain inhibition also promotes mitochondrial reactive oxygen species (ROS) generation in cardiomyocytes,^40–42^ compounding the metabolic injury through oxidative damage to ETC components. The concurrent reduction in the PDH complex, which gates pyruvate flux from glycolysis into the TCA cycle, independently constrains glucose oxidation capacity, together exacerbating the energetic deficit in the failing SAN. To directly assess whether ceramide accumulation is causally sufficient to impair pacemaker cell energetics, we treated iSANCs with mixed ceramides and found that this recapitulated the *in vivo* metabolic phenotype, suppressing fatty acid β-oxidation and driving a glycolytic shift with concomitant ATP depletion (**Fig. 8**). Collectively, these findings are consistent with an emerging view of ceramides as active mediators of mitochondrial dysfunction and energetic failure in the diseased heart, rather than passive markers of metabolic stress. Prior studies have linked ceramides to oxidative stress^43^ and mitochondria-dependent apoptosis.^44,45^

### Functional Implications of Mitochondrial Dysfunction in SAN Automaticity

While our molecular, metabolic, and structural analyses revealed profound mitochondrial derangements in the failing SAN, we determined that these perturbations translated into functional consequences at the cellular level. Importantly, these findings highlight ceramides as essential modulators of SAN electrophysiological function, further supporting the concept that targeting mitochondrial dysfunction may offer a viable therapeutic strategy to restore SAN automaticity and mitigate SAN dysfunction in HF. Recent work by Vaidya et al. demonstrated that sinus node inflammation in HF can also affect SAN automaticity.^11^ Although their study did not focus on metabolic or mitochondrial changes, their findings suggest that immune or inflammatory cues within the SAN, potentially secondary to systemic HF, may directly impact SAN intrinsic function. Taken together, these data suggest that SAN dysfunction in HF arises from a multifactorial interplay of mitochondrial, metabolic, and inflammatory disturbances, each converging to impair the intrinsic automaticity of the SAN. This underscores the need for integrated therapeutic strategies that preserve SAN function by addressing both mitochondrial integrity and the inflammatory microenvironment.

### Study Limitations

The murine multi-omics analyses measure protein and metabolite abundance in mixed SAN tissue and do not establish metabolic flux. Tissue composition differences and cohort effects limit direct comparisons with ventricular tissues. Sample pooling limits assessment of biological variability between individual animals and precludes sex-specific analyses of the multi-omic datasets. For snRNA-seq, SAN tissues from multiple animals were pooled to generate one sequencing sample per condition. Therefore, individual nuclei represent subsamples rather than independent biological replicates. Consequently, the transcriptional analyses characterize condition-associated patterns but do not permit estimation of animal-to-animal variability or population-level differential expression. Although spontaneous action potential recordings from isolated SANCs directly assess intrinsic pacemaker automaticity, sinoatrial conduction time and sinus node recovery time were not measured. Abnormalities therefore in sinoatrial conduction or post-pacing recovery cannot be determined from the present study. Additionally, iSANCs provide a human cellular model but do not fully recapitulate the maturity and physiological context of the adult SAN or establish that endogenous ceramide accumulation is causal. Finally, although the commercial ceramide preparation comprises endogenous long-chain ceramides, the identities and relative proportions of the individual ceramide species are not specified. The observed functional effects therefore cannot be attributed to a specific endogenous ceramide species.

### Conclusions

In the current study, we employed an integrated approach combining multi-omics, ultra-resolution imaging, mitochondrial functional analyses, and electrophysiological assessment to characterize metabolic remodeling associated with SAND in HF. Multi-omic analyses of the HF SAN revealed altered abundance of proteins and metabolites associated with fatty acid β-oxidation, carbohydrate and ketone metabolism, and mitochondrial energy production. Notably, glucosylceramides and ceramides were markedly increased in the HF SAN. In human iSANCs, exogenous mixed ceramide exposure reduced stimulated mitochondrial respiration and ATP-linked energetics, increased extracellular acidification, and impaired spontaneous pacemaker activity in a dose-dependent manner. Ultra-resolution imaging further demonstrated disrupted mitochondrial cristae architecture and reduced structurally modeled ATP-generating capacity in the HF SAN. Together, these findings identify ceramide accumulation as a candidate contributor to metabolic and pacemaker dysfunction in HF and provide a rationale for further investigation of ceramide metabolism as a potential therapeutic target for SAND.

## ACKNOWLEDGEMENTS

This work was supported by the AHA Postdoctoral Fellowship 24POST1198670 (LR); AHA 23CDA1050577 and Tobacco-related Disease Research Program (HZ); NIH F31 HL168956 (DAD); F32HL173968 (AC); NIH R01 DC015135, NIH P01 AG051443, NIH R01 DC01525, NIH R01 DC016099, and NIH R01 AG060504 (ENY); R01 HL171102, R01 HL113006, R01 HL141371, R01 HL141851, R01 HL150693, 2R01HL130020-09A1 (JCW); NIH R01 HL085727, NIH R01 HL085844, NIH R01 HL137228, NIH U01 HL160274, and NIH S10 RR033106, and VA Merit Review Grants I01 BX000576 and I01 CX001490 (NC); NIH T32 HL86350 and NIH F32 HL149288, and Harold S. Geneen Charitable Trust Awards Program for Coronary Heart Disease, AHA Career Development Award 24CDA1276831 (PNT). The ThermoFisher LC/MS system is supported by NIH grant 1S10OD026918-01A1.

## AUTHOR CONTRIBUTIONS

Conceptualization, L.R. N.C. and P.N.T.; methodology, L.R., Y.L., H.Z., N.C., P.N.T., investigation, L.R., Y.L., H.Z., D.A.D., Y.Z., A.C., R.L.W., H.S.S., R.Q.N., G.A.P., G.G., P.N.T; writing – original draft, L.R., P.N.T.; writing – review and editing, L.R., H.Z., N.C. and P.N.T.; funding acquisition, E.N.Y., J.C.W., N.C., P.N.T.; resources, E.N.Y., J.C.W., N.C., P.N.T.; supervision, N.C. and P.N.T.

## SUPPLEMENTARY DATA

### MATERIALS AND METHODS

#### Animal Model

Animal studies were conducted following the approved protocols of the Institutional Animal Care and Use Committee at the University of California, Davis, and adhered to the guidelines published by the US National Institutes of Health (8^th^ Edition). Male and female 10-16-week-old wild-type (WT) C57BL/6J mice (Jackson Laboratory, Bar Harbor, ME) were randomized to undergo sham or transverse aortic constriction (TAC) surgeries and were followed for 8 weeks. TAC surgery was performed as previously described^13^. Briefly, the transverse aorta was visualized and ligated to the size of a 27-gauge needle. The sham procedure was identical, except for the ligation.

#### Echocardiography

We used the VisualSonics Vevo 2100 ultrasound system with the MS 550D probe (FUJIFILM VisualSonics, Inc., Toronto, Canada) to perform echocardiography. Systolic function (B-mode videos and M-mode images) was quantified in conscious mice. Analysis was performed in a blinded fashion.

#### SAN Cell (SANC) Isolation

SANCs were isolated as described previously^46^. Mice were anesthetized by intraperitoneal injection of 80 mg/kg of ketamine and 5 mg/kg of xylazine. Hearts were excised and placed in Tyrode’s solution (35°C) containing (in mM): 140 NaCl, 5.0 HEPES, 5.5 Glucose, 5.4 KCl, 1.8 CaCl_2,_ and 1.0 MgCl_2_ (pH 7.4). The SAN tissue was isolated with a dissecting scope according to the landmarks defined by the orifice of the superior vena cava, crista terminalis, and atrial septum. The SAN tissue was digested in a low Ca2+ solution (pH 6.9) containing collagenase B (0.54 U/mL, Sigma-Aldrich, Burlington, MA), elastase (18.9 U/mL, Sigma), and protease type XIV (1.79 U/mL, Sigma) for 30 minutes at 37°C. After digestion, the tissue was washed with Kraft-Bruhe medium containing (in mM): 100 potassium glutamate, 5 HEPES, 20 glucose, 25 KCl, 10 potassium aspartate, 2 MgSO_4_, 10 KH_2_PO_4_, 20 taurine, 5 creatine, 0.5 EGTA, and 1 mg/mL bovine serum albumin (BSA) (pH 7.4) three times, then SANCs were dissociated with a transfer pipette by mechanically stirring and pipetting the tissue chunks. Dissociated SANCs were stored at room temperature for study.

#### Electrophysiology

SANCs were isolated from sham and TAC mice. Spontaneous single-cell action potentials (APs) were recorded in current-clamp mode using the perforated patch-clamp techniques at 36 ± 0.5 °C^47^. Amphotericin B (240 μg/ml) was added to the pipette solution to form the perforated patch to record APs, using an Axopatch 200A amplifier and a Digidata 1440 digitizer (Molecular Devices, LLC., Sunnyvale, CA). The signals were filtered at 2 kHz using a 4-pole Bessel filter and digitized at a sampling frequency of 5 kHz. All experiments were performed using 3 M KCl agar bridges, connecting the ground electrode to the recording chamber. Borosilicate glass electrodes were pulled with a P-97 micropipette puller (Sutter Instruments, Novato, CA). The resistance of the electrodes was ∼2-3 MΩ when filled with the pipette solutions. Spontaneous APs were recorded in Tyrode’s solution with the pipette filled with (in mM): 30 potassium aspartate, 10 NaCl, 10 HEPES, 0.04 CaCl_2_, 2.0 Mg-ATP, 7.0 phosphocreatine, 0.1 Na-GTP, with pH adjusted to 7.2 with KOH. Data acquisition and analysis were performed using pClamp 10 software (Molecular Devices) and Origin software (OriginLab, Northampton, MA).

#### Transmission Electron Microscopy

Once the hearts were harvested, the SAN tissues were isolated and placed in a fixative containing 2% paraformaldehyde and 2.5% glutaraldehyde in 0.1 M sodium phosphate buffer **o**vernight. The tissues were then rinsed in 0.1M sodium phosphate buffer twice at 15-minute intervals, followed by 45 45-minute incubation in 1% osmium tetroxide. After incubation, the tissues were rinsed with ddH_2_O twice at 15-minute intervals. They then underwent a series of ethanol-induced dehydration, followed by propylene oxide. Pre-infiltration was performed overnight in half propylene oxide, half Poly/Bed Luft resin (Poly/Bed 812, dodecyl succinic anhydride, nadic methyl anhydride, DMP-30). Infiltration was done for 5 hours in 100% Poly/Bed Luft resin. We then embedded the tissues in fresh Poly/Bed Luft resin and allowed the resin to polymerize for 24 hours at 60 °C. Sections of the tissues were taken at approximately 100 nm and collected on copper grids. Sections were stained with 4% aqueous uranyl acetate, rinsed in water, and then treated with 0.3% lead citrate in 0.1 N sodium hydroxide before a final rinse in water. FEI Talos L120C was used to image the sections. Data analysis was performed in a blinded fashion using ImageJ as described.^48^

#### Electron Microscope Tomography

SAN tissues from sham and TAC mice were fixed with 2.5% glutaraldehyde and 2% paraformaldehyde in 0.1 M sodium cacodylate at 37 °C. The tissues were rinsed on ice with 0.1 M sodium cacodylate buffer containing 3 μM CaCl_2_ and incubated in a mixture of 1% OsO4, 0.8% potassium ferrocyanide (C₆FeK₄N₆), and 3 μM CaCl_2_ in 0.1 M sodium cacodylate for 1 hour on ice. Then, the tissues were washed with ice-cold double-distilled water and stained with 2% uranyl acetate for 1 hour on ice, followed by incubation in increasing ethanol solutions (20%, 50%, 70%, 90% on ice, and 3 × 100% at room temperature). Subsequently, the tissues were infiltrated with a mixture of 50% ethanol and 50% Durcupan ACM resin (Fluka) for 6 hours with agitation, and then incubated for 3 × 6-12 hours in 100% Durcupan with agitation. The samples were then polymerized for 48 hours at 60 °C in an oven. Semi-thin sections of approximately 400 nm in thickness were cut from the tissue blocks using a Leica ultramicrotome and placed on 200-mesh, uncoated, thin-bar copper grids. 20-nm colloidal gold particles were deposited on each side of the grid to serve as fiducial cues. The specimens were irradiated for approximately 15 minutes before initiating a double-tilt series to minimize anisotropic specimen thinning during image collection. During data collection, the illumination was maintained at near-parallel beam conditions. Tilt series were captured using SerialEM software (University of Colorado, Boulder, CO) on a Tecnai HiBase Titan (FEI; Hillsboro, OR) electron microscope, operated at 300 kV and with a pixel size of 0.81 nm. Images were recorded with a Gatan 4Kx4K CCD camera. Each double-tilt series consisted of first collecting 121 images taken at 1-degree increments over a range of -60 to +60 degrees, followed by rotating the grid 90 degrees and collecting another 121 images with the same tilt increment. To improve the signal-to-noise ratio, 2x binning was performed by averaging a 2×2 x-y pixel box into 1 pixel.

The IMOD software package (University of Colorado, Boulder, CO) was used for alignment, reconstruction, and volume segmentation. R-weighted back projection was used to generate the reconstructions. Fifty-three mitochondrial volumes were generated for sham and 38 mitochondrial volumes for TAC. Volume segmentation of mitochondrial membranes was performed using IMOD by tracing in each of the 1.62 nm-thick x-y planes in which the object appeared, and then creating stacks of contours with the Drawing Tools plug-in in IMOD. The traced contours were then surface-rendered by turning contours into meshes to generate a 3D model. The surface-rendered volumes were visualized using 3DMOD.

Measurements of mitochondrial outer, inner boundary, and cristae membrane surface areas and volumes were made within segmented volumes using IMODinfo—mitochondrial volume density (defined as the volume occupied by mitochondria divided by the cytoplasmic volume). Number of mitochondria per area, crista density (defined as the total membrane surface area of the cristae divided by the volume of the mitochondrion), and mitochondrion-SR distance were measured using ImageJ.

#### Western Blots

SAN tissues from sham and TAC mice were flash-frozen in liquid nitrogen for western blot experiments. The exact amount of total protein (5 μg) was loaded in each lane. Membranes were blocked in 5% non-fat dry milk (Bio-Rad Laboratories, Hercules, CA) in Tris-buffered saline (TBS) for 1 hour at room temperature and incubated with primary antibodies (1:10000 GAPDH, Abcam, Waltham MA; 1:10000 Total OXPHOS, Abcam) in 5% non-fat dry milk in TBST (TBS containing 0.1% Tween 20) overnight at 4°C. The following day, blots were washed with TBST before incubation with secondary antibodies conjugated with HRP (Abcam). Blots were developed using the ImageQuant LAS 4000 digital imaging system. Quantification was performed with ImageJ.

#### Proteomics (LC/MS)

SAN tissues were isolated exactly as described above. Each sample was a pool of SAN tissues from 3-5 male and female mice. Tissue homogenization was performed as described^49^. Samples were homogenized in a Tris-based Triton X-100 buffer (50 mM Tris-HCl pH 8.5, 5 mM EDTA, 150 mM NaCl, 10 mM KCl, 1% Triton X-100, 5 mM NaF, 5 mM beta-glycerophosphate, 1 mM Na-orthovanadate, containing 1x Roche complete protease inhibitor) in a Dounce Homogenizer on ice. Homogenates were incubated for 2 hrs at 4 °C, sonicated for five cycles (30 s on/30 s off at low amplitude) in a Bioruptor sonicator (Diagenode, USA), cleared by centrifugation (15,000 × g, 20 mins, 4 °C), and supernatant collected.

Proteins from each lysate were subjected to tryptic digestion via suspension-trap (S-Trap) devices (ProtiFi). Disulfide bonds were reduced with dithiothreitol and alkylated with iodoacetamide in 50mM TEAB buffer. The enzymatic digestion involved the initial addition of trypsin at a 1:100 enzyme: protein (wt/wt) ratio for 4 hours at 37 °C, followed by a second addition of trypsin using the same wt/wt ratio for overnight digestion at 37 °C. Peptides were eluted from S-Trap by sequential elution buffers of 100mM TEAB, 0.5% formic acid, and 50% acetonitrile, 0.1% formic acid. The eluted tryptic peptides were dried in a vacuum centrifuge and reconstituted in 0.1% trifluoroacetic acid. These were subjected to LC-MS analysis.

Peptides were resolved on a Thermo Scientific Dionex UltiMate 3000 RSLC system (Waltham MA) using a PepSep (PepSep, Denmark) analytical column: 150umx25cm C18 column, with 1.5 μm particle size (100 Å pores), heated to a constant temperature of 40 °C. Each injection is 0.6 μg of total peptide. Separation was performed in a total run time of 90 mins with a flow rate of 500 μL/min with mobile phases A: water/0.1% formic acid and B: 80%ACN/0.1% formic acid. Peptides were directly eluted onto an Orbitrap Exploris 480 instrument (Thermo Fisher Scientific, Bremen, Germany). Spray voltage was set to 1.8 kV, funnel RF level at 45, and heated capillary temperature at 275 °C. Experiments with full MS resolutions were set to 120,000 at m/z 200, and the full MS AGC target was 300% with an IT of 45 msec. Mass range was set to 350–1200. AGC target value for fragment spectra was set at 1000%. 49 windows of 45.7 Da were used with an overlap of 1 Da. The resolution was set to 30,000 and the IT to Auto. Normalized collision energy was set at 30%. All data were acquired in profile mode using positive polarity, and peptide match was set to off, and isotope exclusion was on.

#### Proteomics Data Analysis

Raw files were processed with Spectronaut version 16 (Biognosys, Zurich, Switzerland) using DirectDIA analysis mode. Mass tolerance/accuracy for precursor and fragment identification was set to the default settings. The Uniprot database of *Mus Musculus* proteins (UP000000589) and a database of 112 common laboratory contaminants (https://www.thegpm.org/crap/) were used. A maximum of two missing cleavages were allowed, the required minimum peptide sequence length was 7 amino acids, and the peptide mass was limited to a maximum of 4600 Da. Carbamidomethylation of cysteine residues was set as a fixed modification, and methionine oxidation and acetylation of protein N termini as variable modifications. A decoy false discovery rate (FDR) at less than 1% for peptide spectrum matches and protein group identifications was used for spectra filtering (Spectronaut default). Decoy database hits, proteins identified as potential contaminants, and proteins identified exclusively by one site modification were excluded from further analysis. For quantification in DIA, we used the standard settings in Spectronaut (version 13). Proteins, which were not quantified in all three replicates for at least one tissue, were discarded for further analysis.

#### Single-nuclei RNA sequencing (snRNA-seq)

SAN tissues were isolated from 7 sham mice and 7 TAC mice, pooled together into two samples, respectively, then snap-frozen in liquid nitrogen and stored at −80 °C until processing. Because tissues were pooled to generate one sequencing sample per condition, individual nuclei were treated as subsamples rather than independent biological replicates. Accordingly, cell-type-resolved transcriptional analyses were used to characterize condition-associated transcriptional patterns and were not interpreted as population-level estimates of biological variability between animals. For nuclei isolation, the two pooled sham and TAC samples were pre-chilled in 5 mL tube on dry ice prior to the addition of 1 mL cold dissociation buffer (5 mM CaCl_2_, 3 mM MgAc, 2 mM EDTA (pH 8.0), 0.5 mM EGTA (pH 8.0), 10 mM Tris-HCl (pH 8.0), 0.1% Triton X-100, 1 mM DTT, 1U/ µL RNase Inhibitor, and RNase-Free Water). The tissues were minced with sterile Iris Scissors in dissociation buffer and incubated for 3 mins on ice. After incubation, the tissue was centrifuged at 40 rcf for 2 mins at 4°C to pellet the large tissue fragments. Next, the supernatant was collected and passed through a 40 µL Flowmi filter into a new pre-chilled 5 mL tube on ice. The filtered supernatant was centrifuged at 1000 rpm for 5 mins at 4°C. After carefully removing the supernatant, the nuclear pellet was resuspended in 500 µL resuspension buffer (1% BSA containing 1 mM DTT and 1 U/µL RNase Inhibitor) and centrifuged again at 1000 rpm for 5 minutes at 4°C. The supernatant was carefully removed, and the nuclei pellet was resuspended in 30 µL resuspension buffer for visual inspection and counting using Trypan Blue. The desired number of nuclei was immediately used to generate single-nucleus libraries according to the 10X protocol (Chromium Next GEM Single Cell 3’ Reagent Kits v3.1 (Dual Index)). Sequencing was performed on an Illumina NovaSeq 6000 using 150PE paired-end reads, yielding approximately 900 million mapped reads per sample.

#### snRNA-seq Data Pre-Processing

For pre-processing of the single-nucleus RNA-seq data, we utilized Chromium Single Cell Software Suite v1.1 from 10X Genomics (Pleasanton, CA). This software suite facilitated sample de-multiplexing, barcode processing, and single-cell 3’ gene counting. Sample de-multiplexing was performed based on the 8-bp sample index read from the Illumina BCL output folder. Subsequently, FASTQ files were generated for the single end reads and sample index using Illumina bcl2fastq. The reads were then mapped to the mouse reference genome (mm10) using STAR. Confidently mapped reads were delivered in a BAM file. Gene-barcode matrices were generated using the Chromium cellular barcodes, and filtered gene-barcode matrices containing only cellular barcodes in MEX format were used for downstream analysis.

#### Data Analysis of snRNA-seq

Only genes expressed in at least three cells and cells with a detected gene count higher than 200 were retained. Additionally, cells with a high percentage of mitochondrial genes (>5%) were filtered out. The Seurat V3 data integration pipeline was used to batch-correct the data using the canonical correlation analysis (CCA) method. According to a benchmark comparison study conducted by Tran and colleagues^50^, Seurat 3 CCA was identified as one of the top three preferred batch integration techniques for this type of data. The R package SCTransform was used to normalize gene expression for each cell by fitting the Gamma-Poisson generalized linear model^51^. The resulting log-transformed, normalized single-cell expression values were used for visualizations and differential expression tests. Statistically significant principal components were identified through a resampling test and retained for the Uniform Manifold Approximation and Projection (UMAP) analysis. Differential expression analysis among clusters was conducted using a likelihood-ratio test, comparing cells within each cluster against all other cells. Gene A was defined as a biomarker for cluster X if it was detected in at least 25% of cells, had an adjusted p-value less than 0.05, and had a log e fold change of at least 0.25 between cells of cluster X and all other cells. These analyses were performed using the Seurat package v4.0. Differentially expressed genes (DEGs) were analyzed for Gene Ontology (GO) terms and Kyoto Encyclopedia of Genes and Genomes (KEGG) pathways enrichment by using KEGG Orthology Based Annotation System (KOBAS)^52^. A significance threshold of FDR < 0.05 was used during the enrichment analysis to identify significant results. The protein-protein interaction network was analyzed by using the STRING website^53^.

### Metabolomics

#### Primary Metabolism

We extracted samples using the Matyash extraction procedure, which includes methyl tert-butyl ether (MTBE), methanol (MeOH), and H_2_O. The organic (upper) phase was dried and resuspended for injection onto the LC, while the aqueous (bottom) phase was dried and submitted for derivatization for gas chromatography (GC). They were shaken at 30°C for 1.5 hrs. We then added 91 µL of *N*-Trimethylsilyl-*N*-methyl trifluoroacetamide (MSTFA) + fatty acid methyl esters (FAMEs) to each sample. Samples were shaken at 37°C for 0.5 hrs to complete derivatization. Samples were then vialed, capped, and injected into the instrument. We use a 7890A GC coupled with a LECO TOF. 0.5 µL of derivatized sample was injected using a splitless method onto a RESTEK RTX-5SIL MS column with an Intergra-Guard at 275°C with a helium flow of 1 mL/min. The GC oven was set to hold at 50°C for 1 min then ramped to 20°C /min to 330°C and then held for 5 mins. The transfer line was set to 280°C while the EI ion source was set to 250°C. The mass spectrometer collected data from 85m/z to 500m/z at an acquisition rate of 17 spectra/sec.

#### Lipidomics

We extracted samples using Matyash extraction procedure which includes MTBE, MeOH, and H_2_O. The organic (upper) phase was dried down and submitted for resuspension and injection onto the LC while the aqueous (bottom) phase was dried down and submitted to derivatization for GC. They are resuspended with 110 uL of a solution of 9:1 methanol: toluene and 50 ng/mL CUDA. This is then shaken for 20 seconds, sonicated for 5 minutes at room temperature, and then centrifuged for 2 minutes at 16100 rcf. The samples are then aliquoted into three parts. 33 uL are aliquoted into a vial with a 50 uL glass insert for positive and negative mode lipidomics. The last part is aliquoted into an Eppendorf tube to be used as a pool. The samples were then loaded on an Agilent 1290 Infinity LC stack. The positive mode was run on an Agilent 6530 with a scan range of m/z 120-1200 Da, with an acquisition speed of 2 spectra/s. A positive mode has between 0.5 and 2 μL injected onto an Acquity Premier BEH C18 1.7 μm, 2.1 × 50 mm Column. The gradient used is 0 min 15% (B), 0.75 min 30% (B), 0.98 min 48% (B), 4.00 min 82% (B), 4.13-4.50 min 99% (B), 4.58-5.50 min 15% (B) with a flow rate of 0.8 mL/min.

The other sample aliquot was run in negative mode, which was run on Agilent 1290 Infinity LC stack, and injected on the same column, with the same gradient and using an Agilent 6546 QTOF mass spec. The acquisition rate was 2 spectra/s with a scan range of m/z 60-1200 Da. The mass resolution for the Agilent 6530 is 10,000 for ESI (+) and 30,000 for ESI (-) for the Agilent 6546.

#### Biogenic Amines

Samples extraction for biogenic amines was performed using a Liquid-Liquid extraction Matyash procedure with MTBE, methanol, and water to create a biphasic partition. The polar phase is then dried down to completeness and run on a Waters Premier Acquity BEH Amide column. A short 4-minute Liquid Chromatography method is used for the separation of polar metabolites from a starting condition of 100% LCMS H2O with 10 mM ammonium formate and 0.125% formic acid to an end condition of 100% ACN:H_2_O 95:5 (v/v) with 10 mM ammonium formate and 0.125% formic acid. A Sciex Triple-ToF scans from 50-1500 m/z with MS/MS collection from 40-1000 selecting from the top 5 ions per cycle. Data processing is performed using MS-Dial, which utilizes an MZ-RT list for annotations in conjunction with a library for MS/MS matching.

#### Differentiation and culture of iSANCs

iPSC-derived SAN cells (iPSC-SANCs) were generated as described in previous publications.^54^ Briefly, human induced pluripotent stem cells were differentiated into SANCs using established protocols that involve stage-specific modulation of signaling pathways. Cells were maintained in SANC maintenance medium under standard culture conditions (37°C, 5% CO₂) until Day 60, before replating for downstream assays.

#### Seahorse fatty acid β oxidation assay

Seahorse fatty acid β-oxidation assay. On day 60 of differentiation, iPSC-SANCs were dissociated using undiluted TrypLE (10×) and seeded into Seahorse XF96 microplates (Agilent Technologies, Santa Clara, CA) at a density of 20,000-40,000 cells per well in RPMI 1640 medium. Cells were maintained under standard culture conditions for 10 days to allow recovery after replating, with medium changes as required. Wells were then randomly assigned to vehicle control or Ceramides Mixture treatment (Selleck Chemicals, E7200; CAS 100403-19-8), which is described by the manufacturer as a mixture of endogenous long-chain ceramides; the individual ceramide species and their relative proportions are not specified by the manufacturer. Ceramides were dissolved in DMSO and administered at final concentrations of 2, 4, or 8 µg/mL. Vehicle control wells received the corresponding concentration of DMSO without ceramides. Treatments were maintained for 14 days, with freshly prepared treatment-containing medium replaced every 2 days.

Fatty acid oxidation was assessed using the Agilent XF Palmitate-BSA FAO Substrate protocol (102720-100). On the day before measurement, growth medium was replaced with substrate-limited medium consisting of DMEM formulated without glucose, glutamine, sodium pyruvate, or HEPES and supplemented with 0.5 mM D-glucose, 1.0 mM GlutaMAX, 0.5 mM L-carnitine, and 1% FBS. Cells were incubated overnight under substrate-limited conditions.

On the assay day, substrate-limited medium was replaced with FAO assay medium prepared from Krebs– Henseleit buffer containing 111 mM NaCl, 4.7 mM KCl, 1.25 mM CaCl_₂_, 2.0 mM MgSO₄, and 1.2 mM NaH₂PO₄, supplemented with 2.5 mM glucose, 0.5 mM L-carnitine, and 5 mM HEPES. The supplemented buffer was prepared on the day of the assay, warmed to 37°C, and adjusted to pH 7.4 with NaOH. Cells were equilibrated in this medium at 37°C in a non-CO₂ incubator for 1 hour before assay initiation.

For FAO inhibition, designated wells were pretreated with 40 µM etomoxir for 15 minutes before recording. Palmitate-BSA was added immediately before measurement according to the manufacturer’s instructions. Baseline oxygen consumption rate (OCR) and extracellular acidification rate (ECAR) were recorded using a Seahorse XF Analyzer. Mitochondrial respiration was subsequently assessed by sequential injection of oligomycin through port A (2.5 µg/mL final), FCCP through port B (2 µM final), and rotenone plus antimycin A through port C (2 and 4 µM final, respectively). OCR and ECAR measurements were normalized to cell number per well and analyzed using Seahorse Wave software.

#### Statistical Analysis

All data are reported as mean±standard error unless otherwise stated. Statistical significance was determined using the student paired t-test. A value of p<0.05 was considered statistically significant.

## FIGURES AND CAPTIONS

**Supplementary Figure 1.**
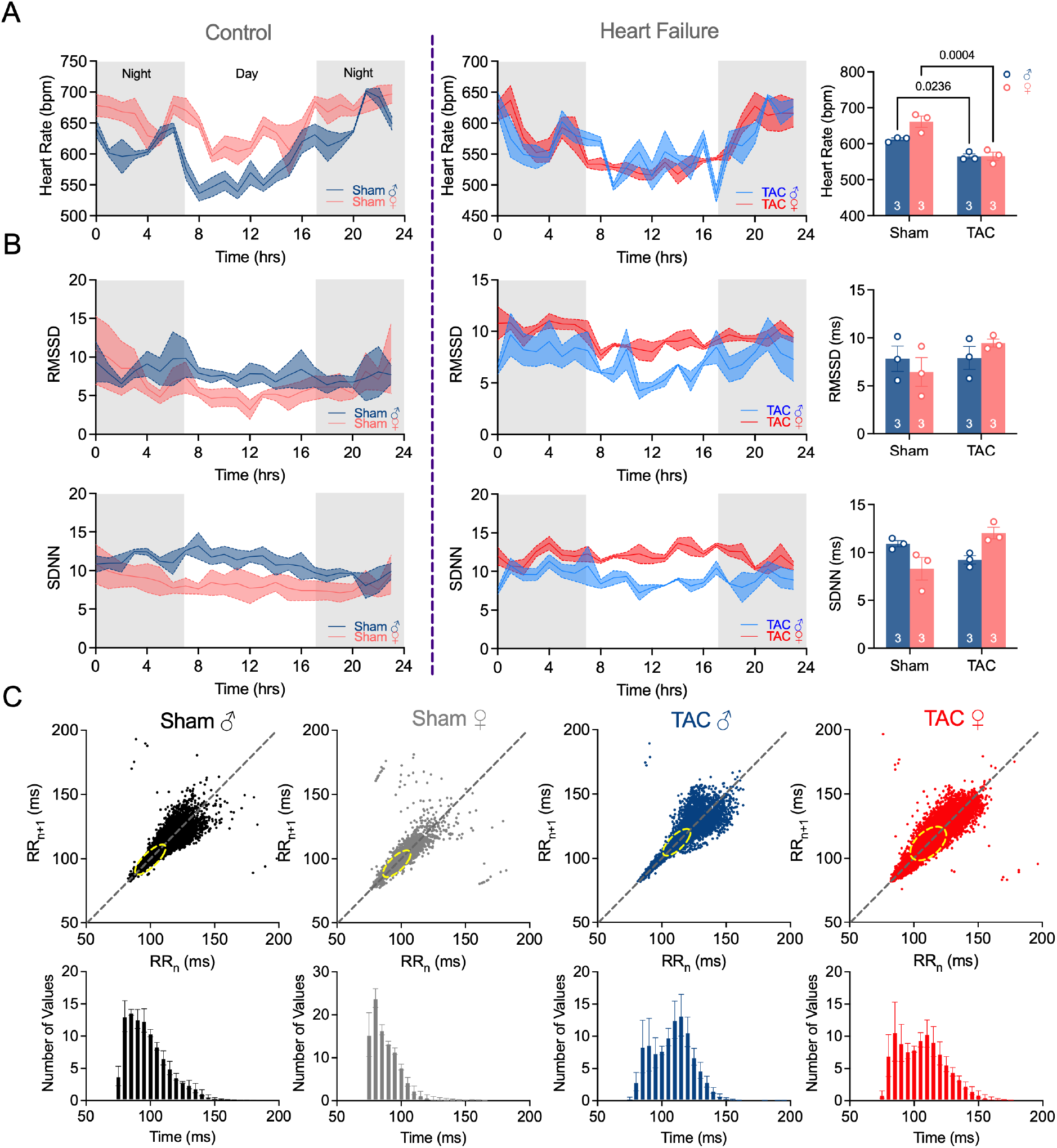
Cardiac electrophysiology and heart rate variability in male and female mice with pressure-overload heart failure. **A)** Twenty-four-hour heart rate telemetry traces in control (Sham) and heart failure (TAC) mice, plotted across light-dark cycles (gray shading = night). Male (blue) and female (pink) mice are shown separately. Bar graphs (right) summarize mean heart rate; TAC significantly reduced heart rate in both sexes. **B)** Heart rate variability assessed by RMSSD (root mean square of successive RR differences, top) and SDNN (standard deviation of normal-to-normal intervals, bottom) over 24 hours in Sham and TAC mice. Summary bar graphs to the right. Neither RMSSD nor SDNN differed significantly between sham and TAC in either sex (n = 3/group). **C)** Poincaré plots (RR_n vs RR_n+1, top) and RR interval frequency histograms (bottom) for each group. The yellow ellipses in the Poincaré plots indicate the central cluster of beat-to-beat intervals. TAC mice show a rightward shift in RR interval distribution consistent with heart rate reduction, without overt loss of short-term variability.

**Supplementary Figure 2.**
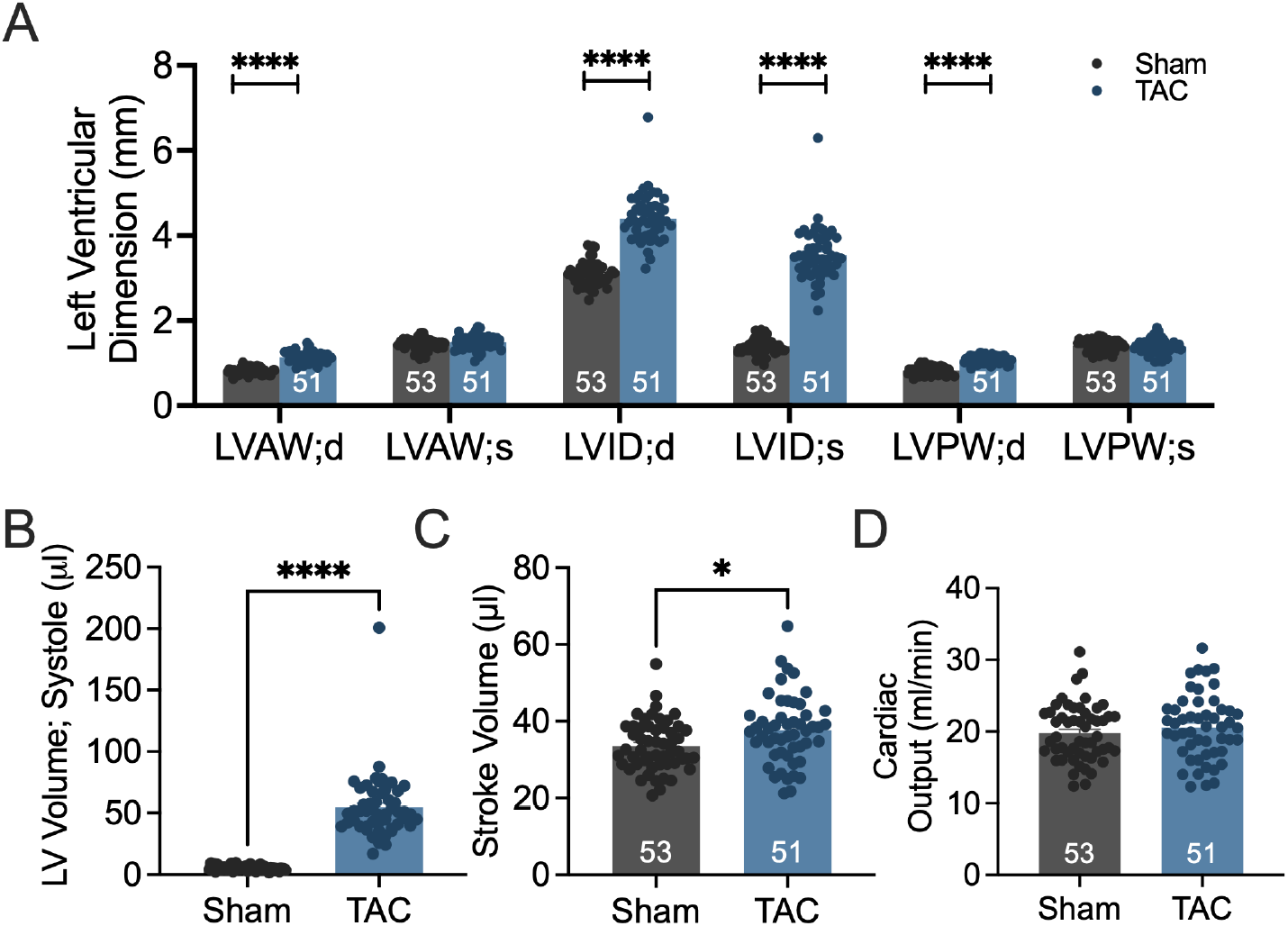
Cardiac structural remodeling in HF. **A**) Summary data of left ventricular dimensions, including left ventricular anterior wall in diastole (LVAW;d), LVAW in systole (LVAW;s), LV inner diameter in diastole (LVID;d), LVID in systole (LVID;s), LV posterior wall in diastole (LVPW;d), and LVPW in systole (LVPW;s). Summary data of **B**) LV volume in systole, **C**) stroke volume, and **D**) cardiac output. Data expressed as mean±SEM. *p<0.05 and ****p<0.0001.

**Supplementary Figure 3.**
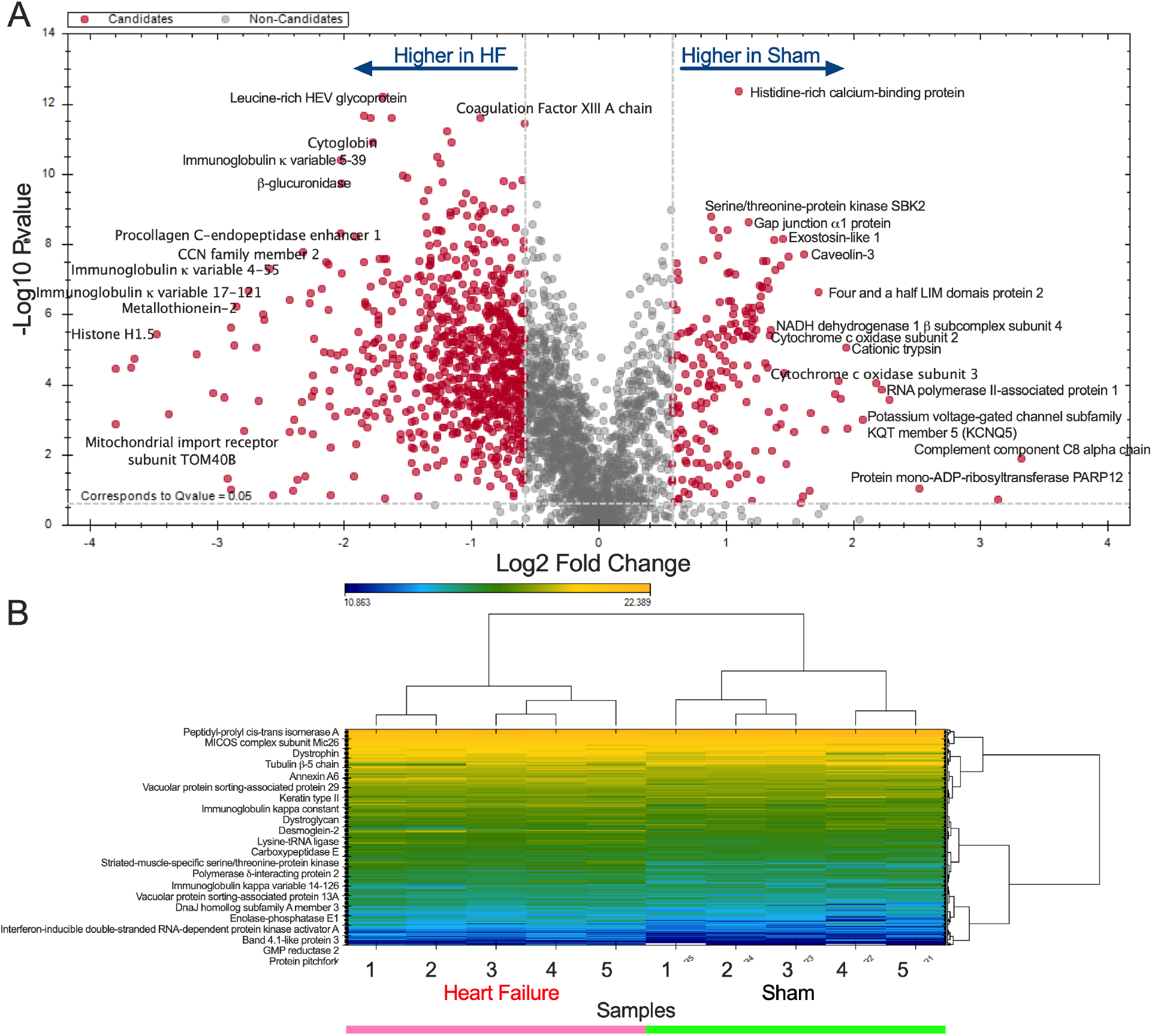
Volcano plot and heat map of all the proteins detected in our proteomics data. **A**) Volcano plot and **B**) heat map illustrating the top differentially expressed proteins between HF and sham groups.

**Supplementary Figure 4.**
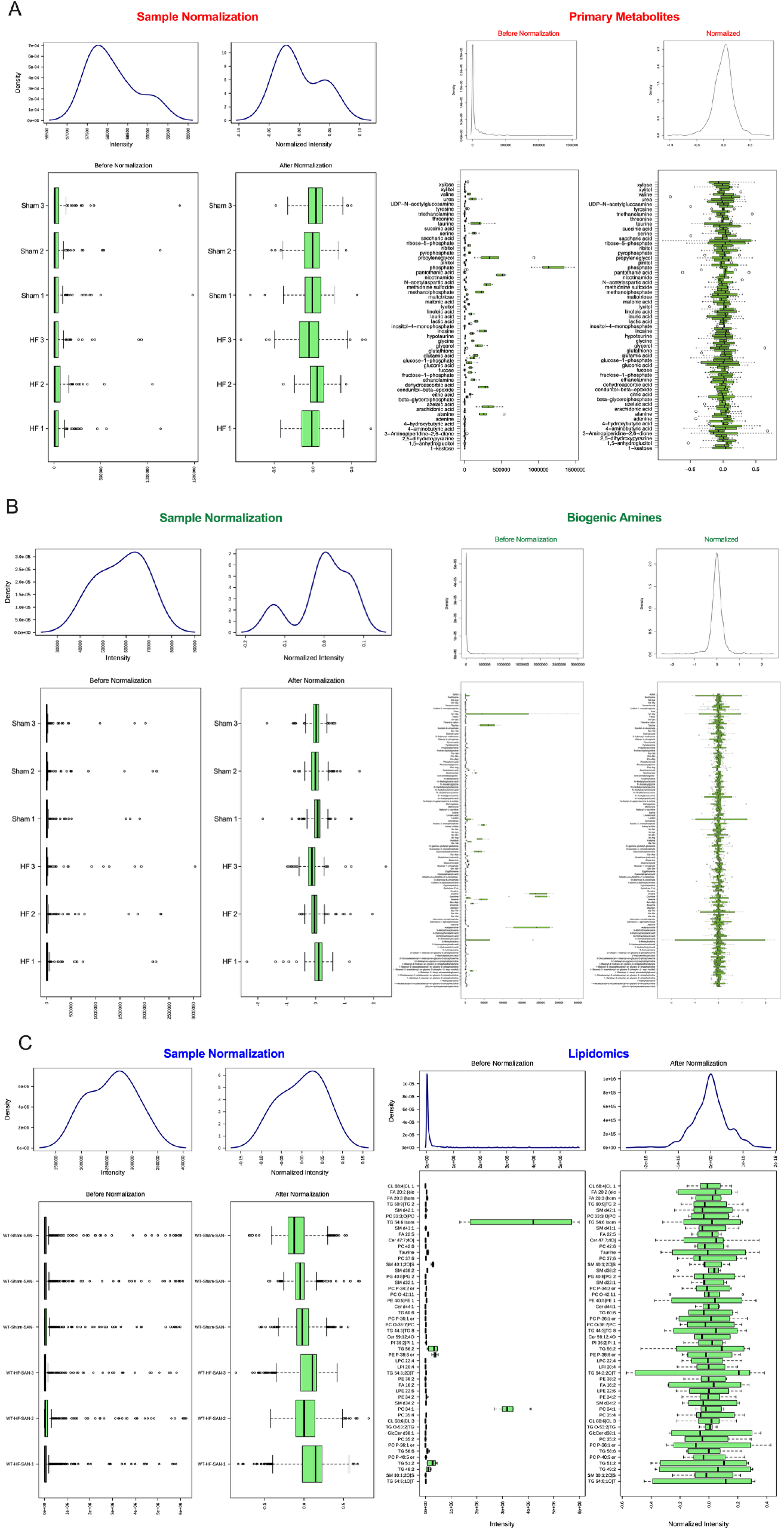
**Normalization of metabolomics data**. **A**) Sample normalization and feature normalization for primary metabolites, **B**) biogenic amines, **C**) and lipidomics to reduce the variation from non-biological sources.

**Supplementary Figure 5.**
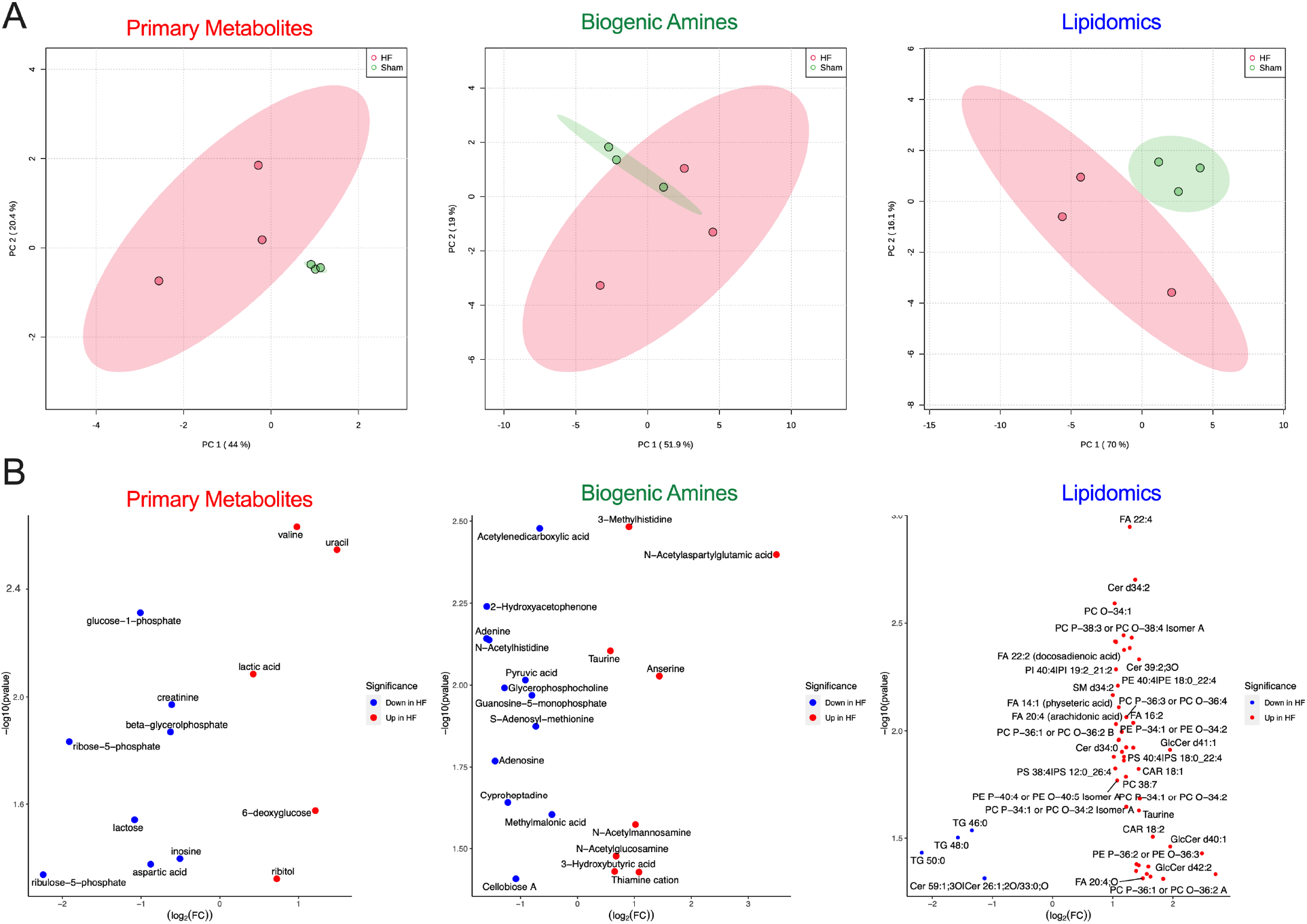
**A**) Principal component analyses and **B**) volcano plots for primary metabolites, biogenic amines, and lipidomics.

**Supplementary Figure 6.**
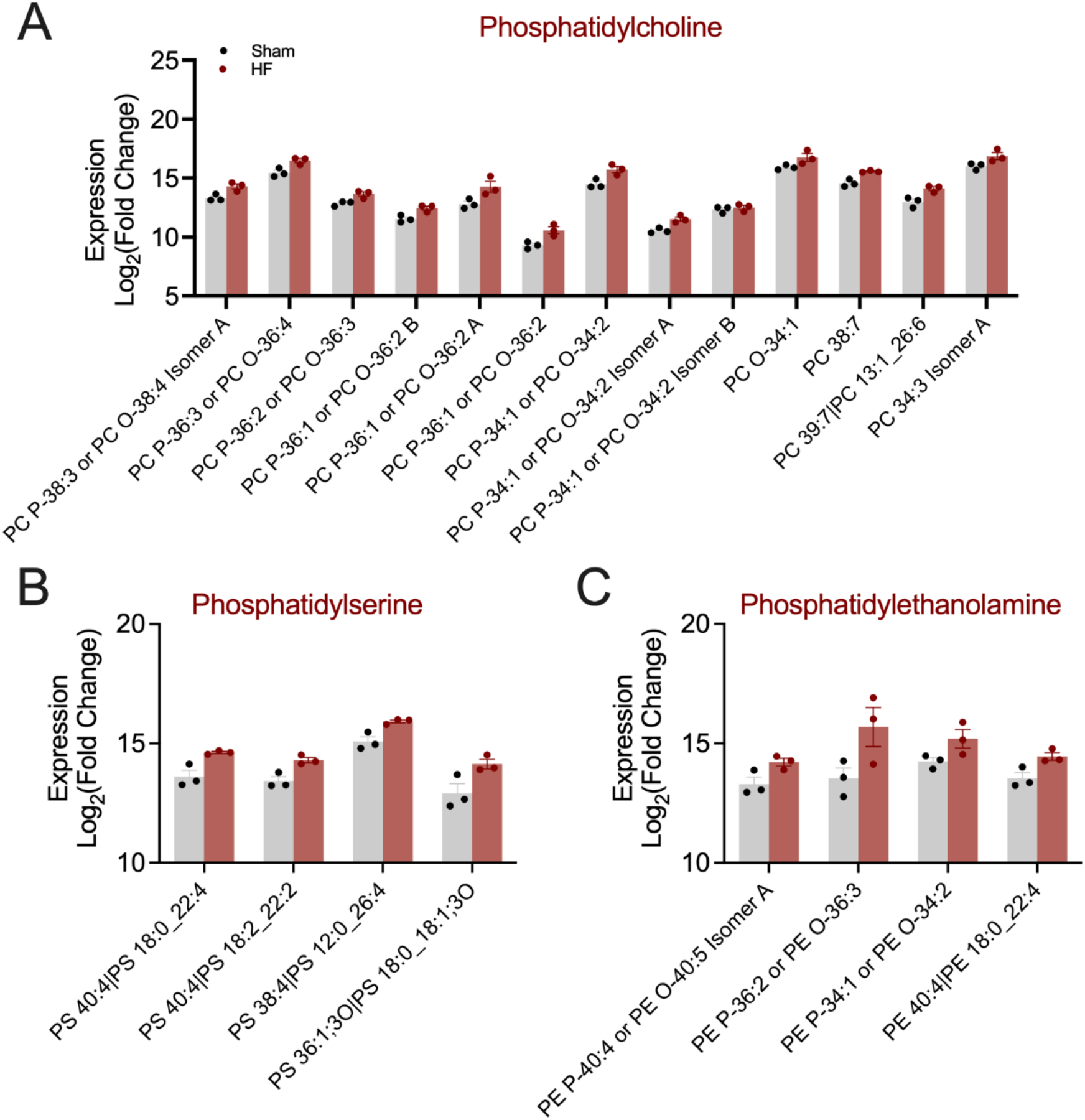
SAN HF exhibited significant alterations in lipid metabolites. Summary data of **A**) phosphatidylcholine, **B**) phosphatidylserine, and **C**) phosphatidylethanolamine.

**Supplementary Figure 7.**
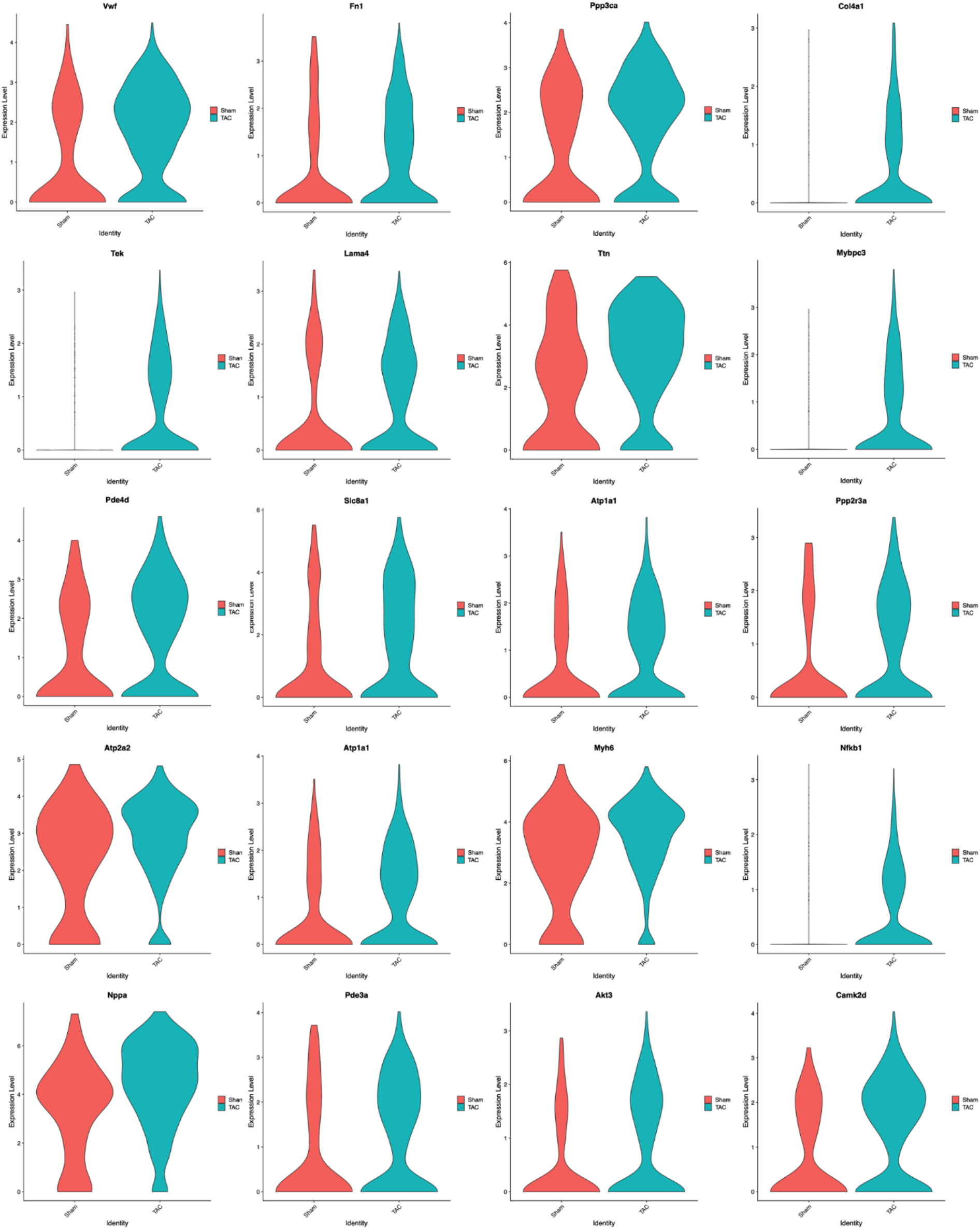
Top 20 differentially expressed metabolic genes between HF and sham groups. Violin plot shows single-nucleus RNA sequencing (snRNA-seq) gene expression levels for the top 20 genes in HF and sham groups. Each panel represents the expression distribution of a specific gene, with HF in blue and sham in red. The width of each violin indicates the density of cells expressing the gene at a given level. Genes shown include markers related to cardiac stress, ion transport, metabolism, and structural remodeling. Differences in gene expression patterns highlight transcriptional alterations associated with HF.

**Supplementary Figure 8.**
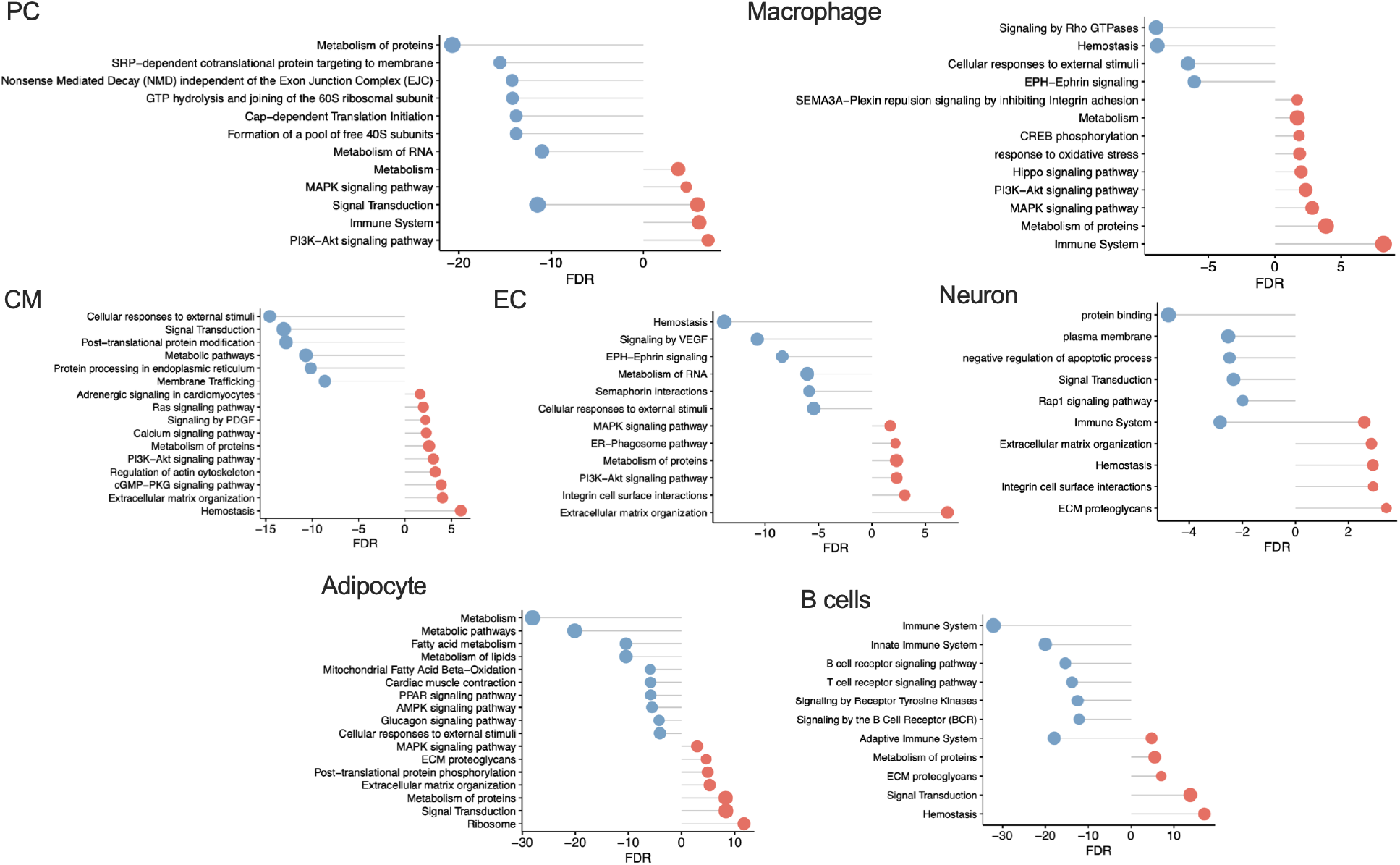
Enriched pathways of different cell types. **A**) Graph shows the enriched pathways of adipocyte, B cells, cardiomyocyte (CM), endothelial cell (EC), fibroblast (FB), macrophage, neuron, and pericyte (PC).

**Supplementary Figure 9.**
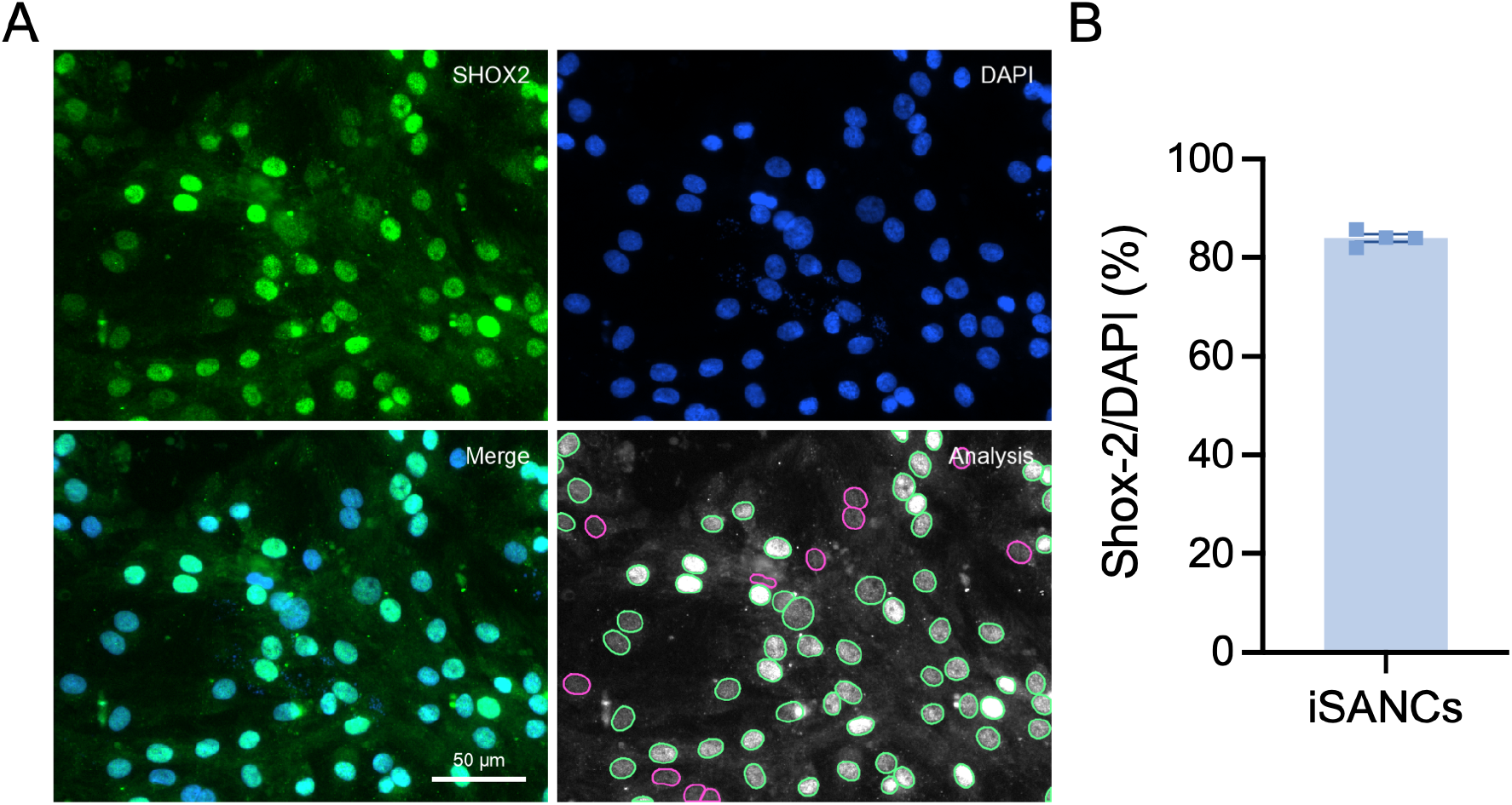
iSAN differentiation efficiency. **A**) Representative fluorescence images of cells stained positive for SHOX2 (SAN marker, green) and DAPI (blue). **B**) Summary graph showing differentiation efficiency.

**Supplementary Figure 10.**
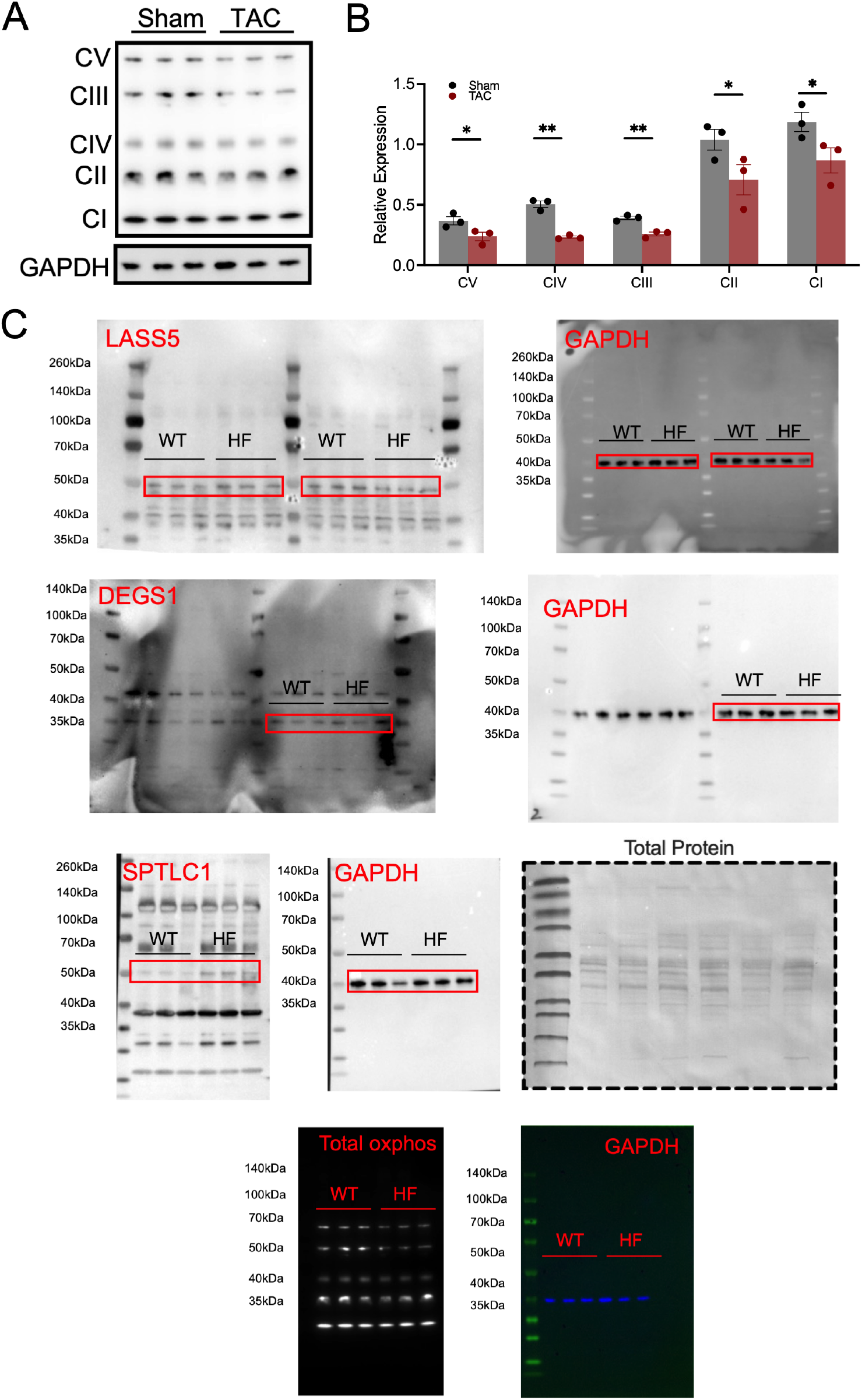
Western blot analyses. **A**) Representative image of Complexes I-V along with **B**) summary data from sham and TAC SAN tissues. **C**) Raw images of all the western blots used in the manuscript. Data expressed as mean±SEM. *p<0.05, **p<0.01.

**Supplementary Table 1.**
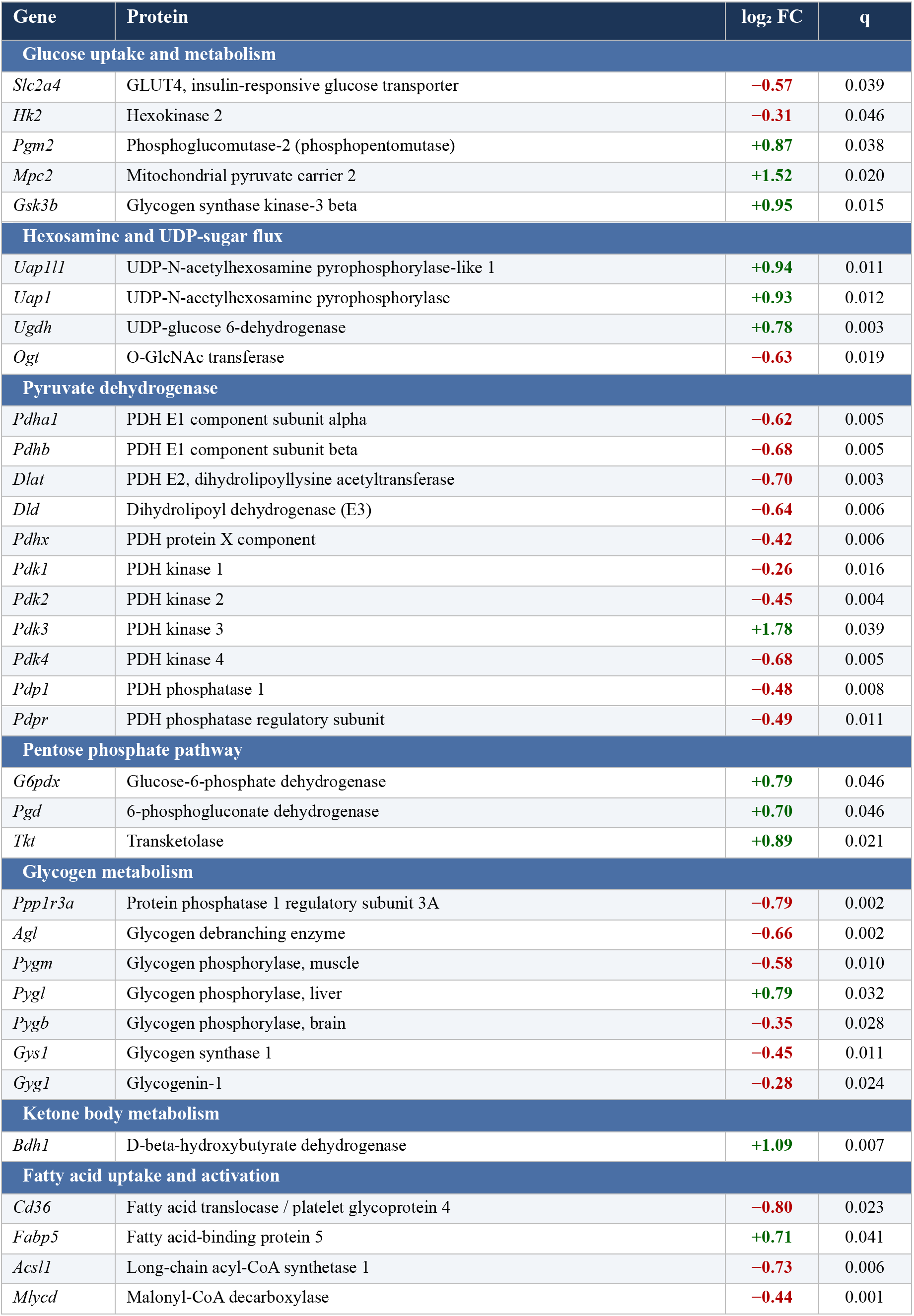
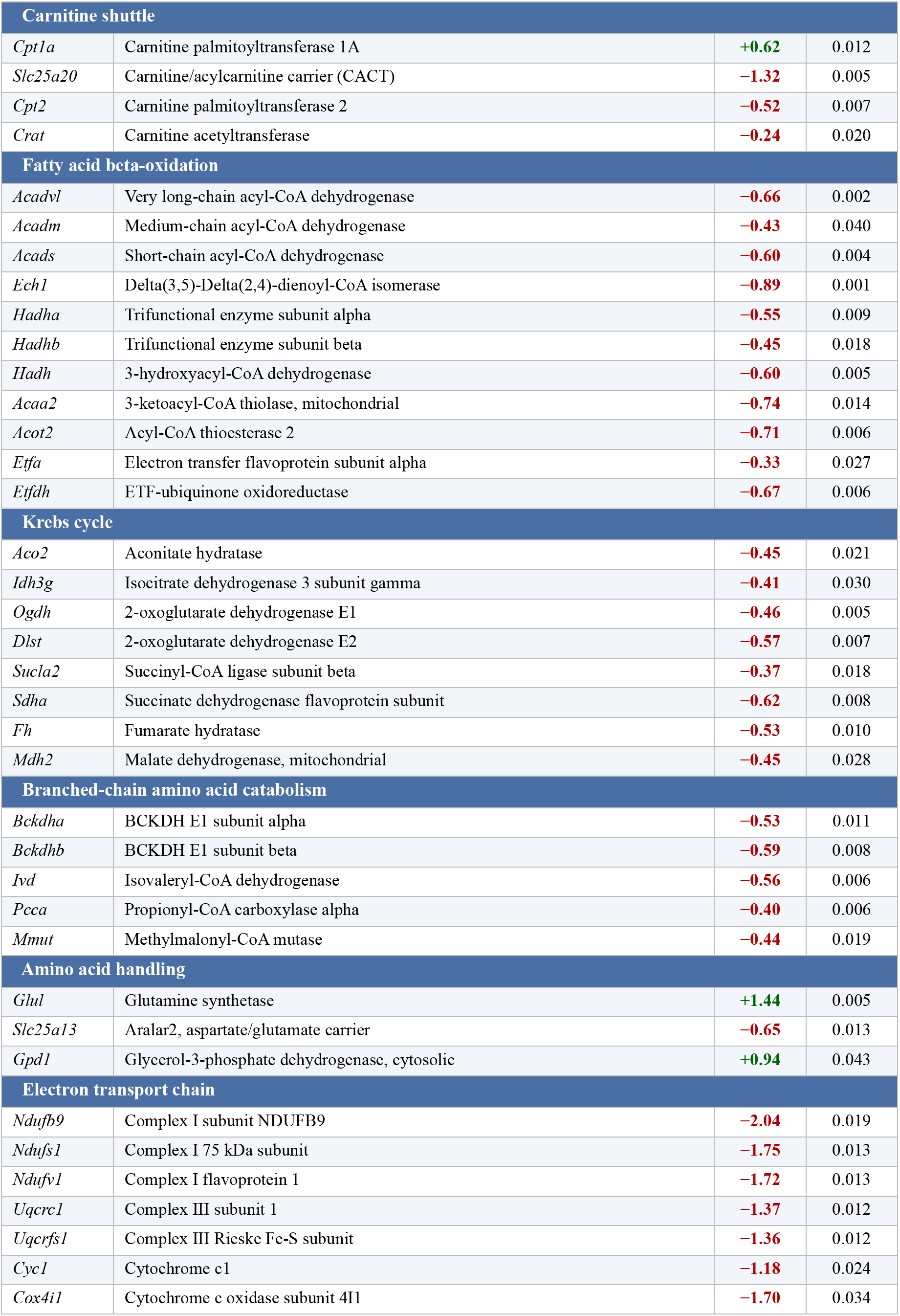
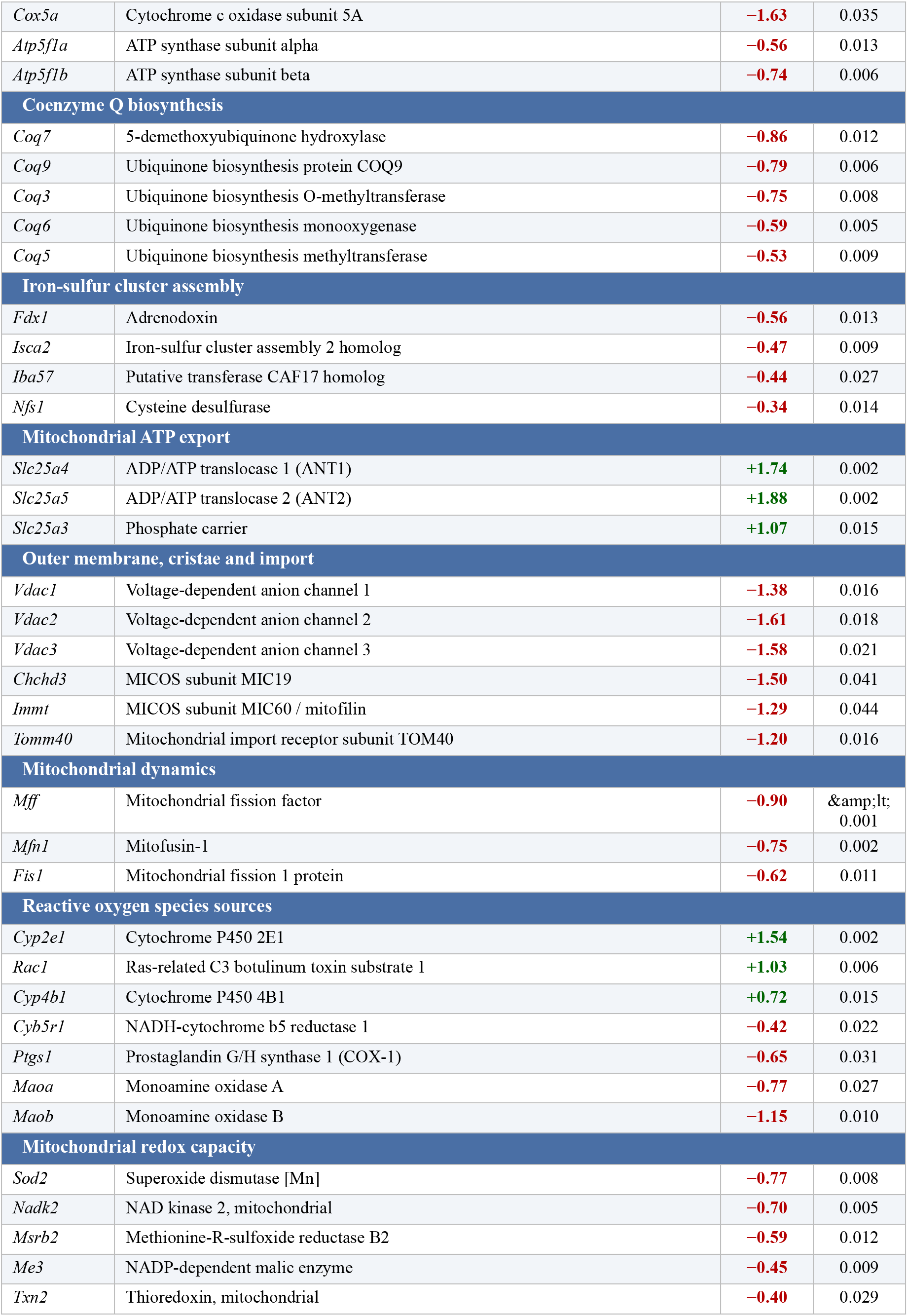
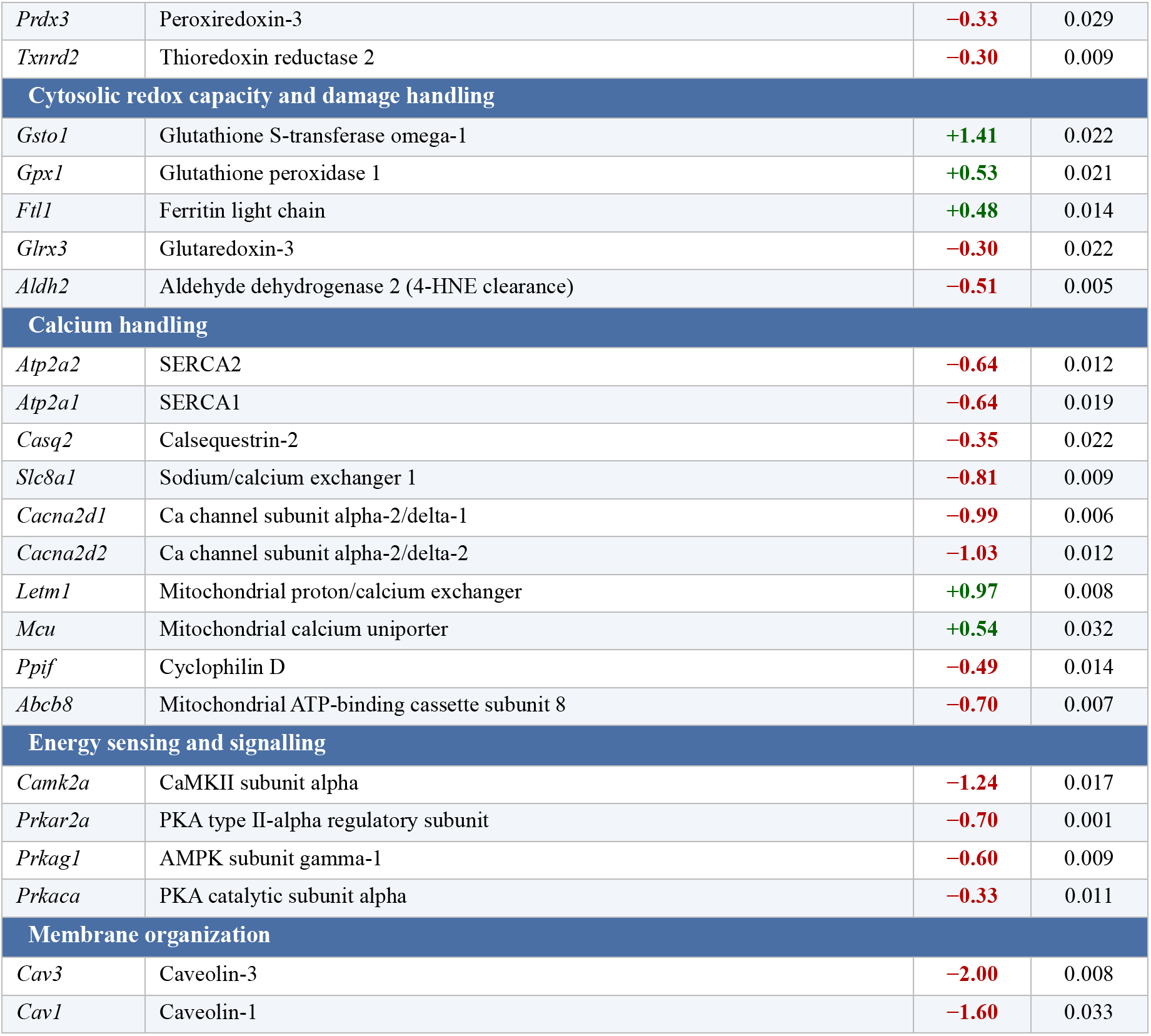
Significant changes in pathways and genes in SANC metabolic pathways. Table shows the pathway, gene name, protein description, log_10_FC, and p-values for the proteins involved in metabolism.

**Supplementary Table 2.** Internal standards used for lipidomics. The table lists the different standards used to ensure quality control for the metabolomics data.

| Lipid Internal Standard | ng/mL | μmol | nmol |
| --- | --- | --- | --- |
| 1_CE(22:1) iSTD | 16919.3 | 23.9 | 23924 |
| 1_PE(17:0/17:0) iSTD | 369.8 | 0.51 | 513.5 |
| 1_PG (17:0/17:0) iSTD | 1479.1 | 1.97 | 1972 |
| 1_LPC(17:0) iSTD | 246.5 | 0.48 | 483.7 |
| 1_Sphingosine(d17:1) iSTD | 54.8 | 0.19 | 191.9 |
| 1_Ceramide (d18:1/17:0) iSTD | 123.3 | 0.22 | 223.3 |
| 1_SM (d18:1/17:0) iSTD | 98.6 | 0.14 | 137.5 |
| 1_FA (16:0)-d3 iSTD | 171.2 | 0.66 | 662.4 |
| 1_PC(12:0/13:0) iSTD | 9.9 | 0.015 | 15.5 |
| 1_Cholesterol d7 iSTD | 493.0 | 1.25 | 1252.3 |
| 1_TAG d5(17:0/17:1/17:0) iSTD | 154.1 | 0.18 | 180.8 |
| 1_DG(12:0/12:0/0:0) iSTD | 493.0 | 1.08 | 1079.6 |
| 1_DG(18:1/2:0/0:0) iSTD | 2958.2 | 7.42 | 7422.1 |
| 1_MG (17:0/0:0/0:0) iSTD | 986.1 | 2.86 | 2862.1 |
| 1_LPE (17:1) iSTD | 123.3 | 0.26 | 264.8 |

**Supplementary Table 3.**
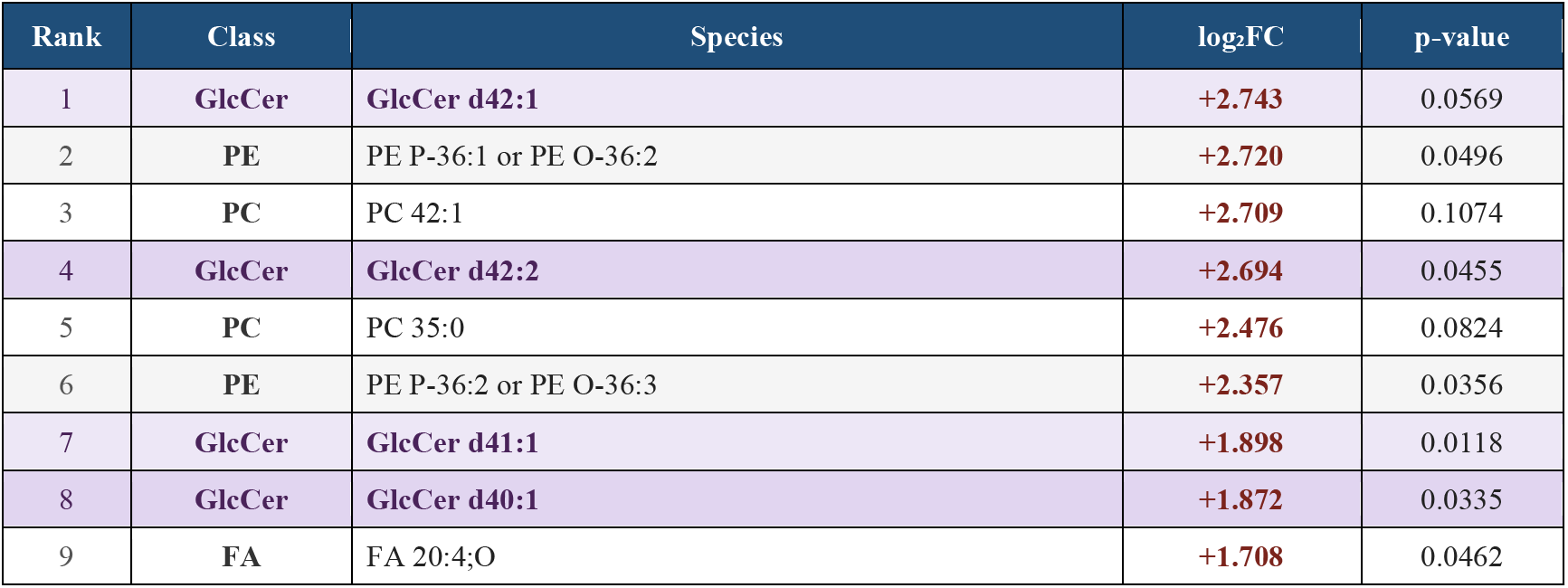
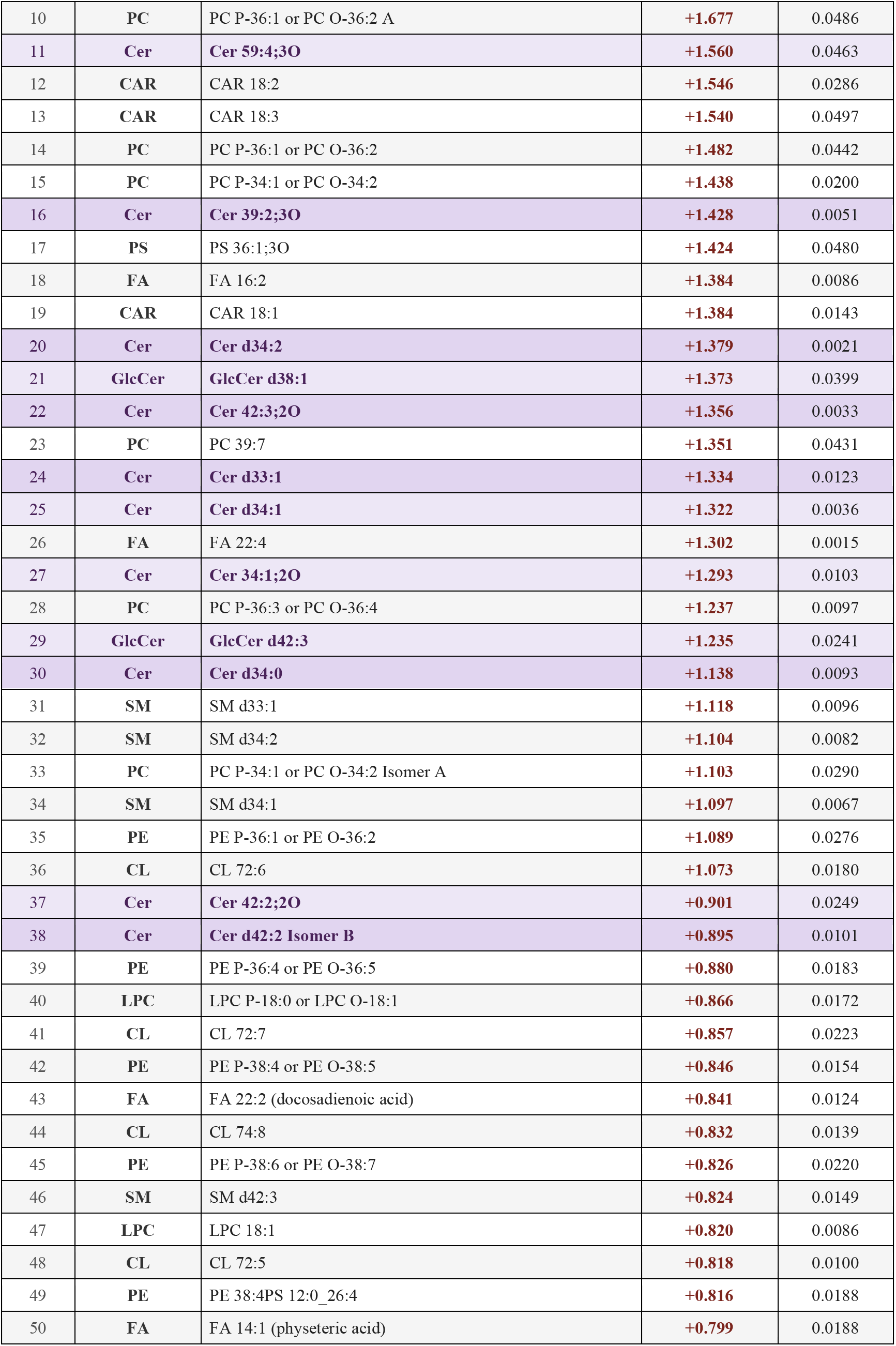
Top upregulated lipids in the SAN in HF. Table shows the rank, class, species, log2FC, p-value, and FDR of upregulated lipids in lipidomics dataset.

| Rank | Class | Species | log<sub>2</sub>FC | p-value |
| --- | --- | --- | --- | --- |
| 1 | GlcCer | GlcCer d42:1 | +2.743 | 0.0569 |
| 2 | PE | PE P-36:1 or PE O-36:2 | +2.720 | 0.0496 |
| 3 | PC | PC 42:1 | +2.709 | 0.1074 |
| 4 | GlcCer | GlcCer d42:2 | +2.694 | 0.0455 |
| 5 | PC | PC 35:0 | +2.476 | 0.0824 |
| 6 | PE | PE P-36:2 or PE O-36:3 | +2.357 | 0.0356 |
| 7 | GlcCer | GlcCer d41:1 | +1.898 | 0.0118 |
| 8 | GlcCer | GlcCer d40:1 | +1.872 | 0.0335 |
| 9 | FA | FA 20:4;O | +1.708 | 0.0462 |
| 10 | PC | PC P-36:1 or PC O-36:2 A | +1.677 | 0.0486 |
| 11 | Cer | Cer 59:4;3O | +1.560 | 0.0463 |
| 12 | CAR | CAR 18:2 | +1.546 | 0.0286 |
| 13 | CAR | CAR 18:3 | +1.540 | 0.0497 |
| 14 | PC | PC P-36:1 or PC O-36:2 | +1.482 | 0.0442 |
| 15 | PC | PC P-34:1 or PC O-34:2 | +1.438 | 0.0200 |
| 16 | Cer | Cer 39:2;3O | +1.428 | 0.0051 |
| 17 | PS | PS 36:1;3O | +1.424 | 0.0480 |
| 18 | FA | FA 16:2 | +1.384 | 0.0086 |
| 19 | CAR | CAR 18:1 | +1.384 | 0.0143 |
| 20 | Cer | Cer d34:2 | +1.379 | 0.0021 |
| 21 | GlcCer | GlcCer d38:1 | +1.373 | 0.0399 |
| 22 | Cer | Cer 42:3;2O | +1.356 | 0.0033 |
| 23 | PC | PC 39:7 | +1.351 | 0.0431 |
| 24 | Cer | Cer d33:1 | +1.334 | 0.0123 |
| 25 | Cer | Cer d34:1 | +1.322 | 0.0036 |
| 26 | FA | FA 22:4 | +1.302 | 0.0015 |
| 27 | Cer | Cer 34:1;2O | +1.293 | 0.0103 |
| 28 | PC | PC P-36:3 or PC O-36:4 | +1.237 | 0.0097 |
| 29 | GlcCer | GlcCer d42:3 | +1.235 | 0.0241 |
| 30 | Cer | Cer d34:0 | +1.138 | 0.0093 |
| 31 | SM | SM d33:1 | +1.118 | 0.0096 |
| 32 | SM | SM d34:2 | +1.104 | 0.0082 |
| 33 | PC | PC P-34:1 or PC O-34:2 Isomer A | +1.103 | 0.0290 |
| 34 | SM | SM d34:1 | +1.097 | 0.0067 |
| 35 | PE | PE P-36:1 or PE O-36:2 | +1.089 | 0.0276 |
| 36 | CL | CL 72:6 | +1.073 | 0.0180 |
| 37 | Cer | Cer 42:2;2O | +0.901 | 0.0249 |
| 38 | Cer | Cer d42:2 Isomer B | +0.895 | 0.0101 |
| 39 | PE | PE P-36:4 or PE O-36:5 | +0.880 | 0.0183 |
| 40 | LPC | LPC P-18:0 or LPC O-18:1 | +0.866 | 0.0172 |
| 41 | CL | CL 72:7 | +0.857 | 0.0223 |
| 42 | PE | PE P-38:4 or PE O-38:5 | +0.846 | 0.0154 |
| 43 | FA | FA 22:2 (docosadienoic acid) | +0.841 | 0.0124 |
| 44 | CL | CL 74:8 | +0.832 | 0.0139 |
| 45 | PE | PE P-38:6 or PE O-38:7 | +0.826 | 0.0220 |
| 46 | SM | SM d42:3 | +0.824 | 0.0149 |
| 47 | LPC | LPC 18:1 | +0.820 | 0.0086 |
| 48 | CL | CL 72:5 | +0.818 | 0.0100 |
| 49 | PE | PE 38:4PS 12:0 26:4 | +0.816 | 0.0188 |
| 50 | FA | FA 14:1 (physeteric acid) | +0.799 | 0.0188 |

**Supplementary Table 4.**
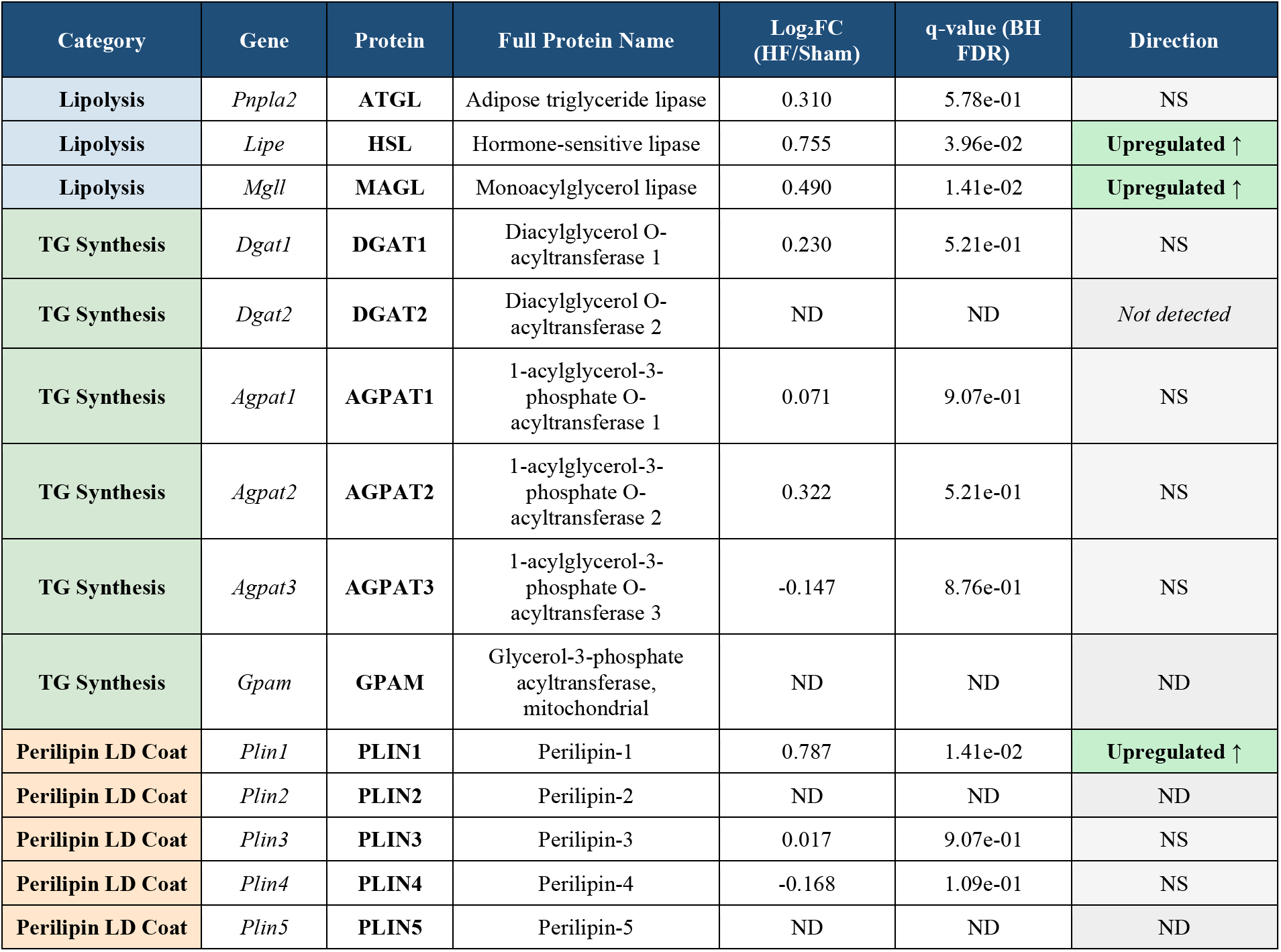
Proteomic abundance of lipid droplet-associated proteins in SAN HF. Log₂FC, log₂ fold change (HF/Sham) calculated from per-sample intensities (n=5/group). q-values from Benjamini–Hochberg FDR correction across detected proteins. Significance: q < 0.05. ND: not detected in proteomic dataset.

## REFERENCES

1 Bui, A. L., Horwich, T. B. & Fonarow, G. C. Epidemiology and risk profile of heart failure. Nat Rev Cardiol 8, 30–41 (2011). 10.1038/nrcardio.2010.165

2 Clare, J. T. et al. Trends in survival after a diagnosis of heart failure in the United Kingdom 2000-2017: population based cohort study. BMJ 364, l223 (2019). 10.1136/bmj.l223

3 Tomaselli, G. F. et al. Sudden cardiac death in heart failure. The role of abnormal repolarization. Circulation 90, 2534–2539 (1994). 10.1161/01.CIR.90.5.2534

4 Nohria, A., Lewis, E. & Stevenson, L. W. Medical Management of Advanced Heart Failure. JAMA 287, 628–640 (2002). 10.1001/jama.287.5.628

5 Mann, D. L. & Bristow, M. R. Mechanisms and Models in Heart Failure. Circulation 111, 2837–2849 (2005). 10.1161/CIRCULATIONAHA.104.500546

6 Lou, Q. et al. Upregulation of adenosine A1 receptors facilitates sinoatrial node dysfunction in chronic canine heart failure by exacerbating nodal conduction abnormalities revealed by novel dual-sided intramural optical mapping. Circulation 130, 315–324 (2014). 10.1161/circulationaha.113.007086

7 Sanders, P., Kistler, P. M., Morton, J. B., Spence, S. J. & Kalman, J. M. Remodeling of Sinus Node Function in Patients With Congestive Heart Failure. Circulation 110, 897–903 (2004). 10.1161/01.CIR.0000139336.69955.AB

8 Mesquita, T. et al. Mechanisms of Sinoatrial Node Dysfunction in Heart Failure With Preserved Ejection Fraction. Circulation 145, 45–60 (2022). 10.1161/CIRCULATIONAHA.121.054976

9 Xue, J. B. et al. Heart failure in mice induces a dysfunction of the sinus node associated with reduced CaMKII signaling. J Gen Physiol 154 (2022). 10.1085/jgp.202112895

10 Opthof, T. et al. Changes in sinus node function in a rabbit model of heart failure with ventricular arrhythmias and sudden death. Circulation 101, 2975–2980 (2000). 10.1161/01.cir.101.25.2975

11 Kahnert, K. et al. Proteomics couples electrical remodelling to inflammation in a murine model of heart failure with sinus node dysfunction. Cardiovascular Research 120, 927–942 (2024). 10.1093/cvr/cvae054

12 Zicha, S., Fernández-Velasco, M., Lonardo, G., L’Heureux, N. & Nattel, S. Sinus node dysfunction and hyperpolarization-activated (HCN) channel subunit remodeling in a canine heart failure model. Cardiovascular Research 66, 472–481 (2005). 10.1016/j.cardiores.2005.02.011

13 Ren, L. et al. Disruption of mitochondria–sarcoplasmic reticulum microdomain connectomics contributes to sinus node dysfunction in heart failure. Proceedings of the National Academy of Sciences 119, e2206708119 (2022). doi:10.1073/pnas.2206708119

14 Ingwall, J. S. & Weiss, R. G. Is the Failing Heart Energy Starved? Circulation Research 95, 135–145 (2004). 10.1161/01.RES.0000137170.41939.d9

15 Lopaschuk, G. D., Karwi, Q. G., Tian, R., Wende, A. R. & Abel, E. D. Cardiac Energy Metabolism in Heart Failure. Circulation Research 128, 1487–1513 (2021). 10.1161/CIRCRESAHA.121.318241

16 Ventura-Clapier, R., Garnier, A., Veksler, V. & Joubert, F. Bioenergetics of the failing heart. Biochimica et Biophysica Acta (BBA) - Molecular Cell Research 1813, 1360–1372 (2011). 10.1016/j.bbamcr.2010.09.006

17 Ren, L. et al. Disruption of mitochondria-sarcoplasmic reticulum microdomain connectomics contributes to sinus node dysfunction in heart failure. Proc Natl Acad Sci U S A 119, e2206708119 (2022). 10.1073/pnas.2206708119

18 Diloretto, D. A. et al. Ro5-4864, a ligand of the mitochondrial translocator protein, protects against heart failure in mice via regulation of the p62-Keap1-Nrf2 axis. J Physiol 604, 6943–6958 (2026). 10.1113/jp288457

19 Afzal, N., Lederer, W. J., Jafri, M. S. & Mannella, C. A. Effect of crista morphology on mitochondrial ATP output: A computational study. Curr Res Physiol 4, 163–176 (2021). 10.1016/j.crphys.2021.03.005

20 Song, D. H. et al. Biophysical significance of the inner mitochondrial membrane structure on the electrochemical potential of mitochondria. Phys Rev E Stat Nonlin Soft Matter Phys 88, 062723 (2013). 10.1103/PhysRevE.88.062723

21 Benador, I. Y., Veliova, M., Liesa, M. & Shirihai, O. S. Mitochondria Bound to Lipid Droplets: Where Mitochondrial Dynamics Regulate Lipid Storage and Utilization. Cell Metab 29, 827–835 (2019). 10.1016/j.cmet.2019.02.011

22 Goldberg, I. J., Trent, C. M. & Schulze, P. C. Lipid metabolism and toxicity in the heart. Cell Metab 15, 805–812 (2012). 10.1016/j.cmet.2012.04.006

23 Son, N. H. et al. Cardiac lipid droplets differ under pathological and physiological conditions. J Lipid Res 66, 100920 (2025). 10.1016/j.jlr.2025.100920

24 Matsuura, T. R., Puchalska, P., Crawford, P. A. & Kelly, D. P. Ketones and the Heart: Metabolic Principles and Therapeutic Implications. Circulation Research 132, 882–898 (2023). 10.1161/CIRCRESAHA.123.321872

25 Lommi, J. et al. Blood ketone bodies in congestive heart failure. J Am Coll Cardiol 28, 665–672 (1996). 10.1016/0735-1097(96)00214-8

26 Doan, M. T. et al. Direct anabolic metabolism of three-carbon propionate to a six-carbon metabolite occurs in&#xa0;vivo across tissues and species. Journal of Lipid Research 63 (2022). 10.1016/j.jlr.2022.100224

27 Taegtmeyer, H., Sen, S. & Vela, D. Return to the fetal gene program: a suggested metabolic link to gene expression in the heart. Ann N Y Acad Sci 1188, 191–198 (2010). 10.1111/j.1749-6632.2009.05100.x

28 van Bilsen, M., Smeets, P. J., Gilde, A. J. & van der Vusse, G. J. Metabolic remodelling of the failing heart: the cardiac burn-out syndrome? Cardiovascular research 61, 218–226 (2004).

29 Peterson, L. R. et al. Ceramide Remodeling and Risk of Cardiovascular Events and Mortality. J Am Heart Assoc 7 (2018). 10.1161/jaha.117.007931

30 Choi, R. H., Tatum, S. M., Symons, J. D., Summers, S. A. & Holland, W. L. Ceramides and other sphingolipids as drivers of cardiovascular disease. Nature reviews. Cardiology 18, 701–711 (2021). 10.1038/s41569-021-00536-1

31 Chaurasia, B. & Summers, S. A. Ceramides – Lipotoxic Inducers of Metabolic Disorders. Trends in Endocrinology & Metabolism 26, 538–550 (2015). 10.1016/j.tem.2015.07.006

32 Fucho, R., Casals, N., Serra, D. & Herrero, L. Ceramides and mitochondrial fatty acid oxidation in obesity. Faseb j 31, 1263–1272 (2017). 10.1096/fj.201601156R

33 Vos, M. et al. Ceramide accumulation induces mitophagy and impairs β-oxidation in PINK1 deficiency. Proceedings of the National Academy of Sciences 118, e2025347118 (2021). 10.1073/pnas.2025347118

34 Di Paola, M., Cocco, T. & Lorusso, M. Ceramide interaction with the respiratory chain of heart mitochondria. Biochemistry 39, 6660–6668 (2000).

35 Law, B. A. et al. Lipotoxic very-long-chain ceramides cause mitochondrial dysfunction, oxidative stress, and cell death in cardiomyocytes. FASEB journal : official publication of the Federation of American Societies for Experimental Biology 32, 1403–1416 (2018). 10.1096/fj.201700300r

36 Schubert, K. M., Scheid, M. P. & Duronio, V. Ceramide Inhibits Protein Kinase B/Akt by Promoting Dephosphorylation of Serine 473*. Journal of Biological Chemistry 275, 13330–13335 (2000). 10.1074/jbc.275.18.13330

37 Summers, S. A., Garza, L. A., Zhou, H. & Birnbaum, M. J. Regulation of insulin-stimulated glucose transporter GLUT4 translocation and Akt kinase activity by ceramide. Mol Cell Biol 18, 5457–5464 (1998). 10.1128/mcb.18.9.5457

38 Diaz-Vegas, A. et al. Mitochondrial electron transport chain, ceramide, and coenzyme Q are linked in a pathway that drives insulin resistance in skeletal muscle. eLife 12, RP87340 (2023). 10.7554/eLife.87340

39 Kogot-Levin, A. & Saada, A. Ceramide and the mitochondrial respiratory chain. Biochimie 100, 88–94 (2014). 10.1016/j.biochi.2013.07.027

40 Di Paola, M., Cocco, T. & Lorusso, M. Ceramide interaction with the respiratory chain of heart mitochondria. Biochemistry 39, 6660–6668 (2000). 10.1021/bi9924415

41 Law, B. A. et al. Lipotoxic very-long-chain ceramides cause mitochondrial dysfunction, oxidative stress, and cell death in cardiomyocytes. Faseb j 32, 1403–1416 (2018). 10.1096/fj.201700300R

42 Bekhite, M. et al. The role of ceramide accumulation in human induced pluripotent stem cell-derived cardiomyocytes on mitochondrial oxidative stress and mitophagy. Free Radic Biol Med 167, 66–80 (2021). 10.1016/j.freeradbiomed.2021.02.016

43 Ding, S. et al. Ceramides and mitochondrial homeostasis. Cell Signal 117, 111099 (2024). 10.1016/j.cellsig.2024.111099

44 Parra, V. et al. Changes in mitochondrial dynamics during ceramide-induced cardiomyocyte early apoptosis. Cardiovasc Res 77, 387–397 (2008). 10.1093/cvr/cvm029

45 Parra, V. et al. Calcium and mitochondrial metabolism in ceramide-induced cardiomyocyte death. Biochim Biophys Acta 1832, 1334–1344 (2013). 10.1016/j.bbadis.2013.04.009

46 Fenske, S. et al. Comprehensive multilevel in vivo and in vitro analysis of heart rate fluctuations in mice by ECG telemetry and electrophysiology. Nat Protoc 11, 61–86 (2016). 10.1038/nprot.2015.139

47 Hamill, O. P., Marty, A., Neher, E., Sakmann, B. & Sigworth, F. J. Improved patch-clamp techniques for high-resolution current recording from cells and cell-free membrane patches. Pflugers Arch 391, 85–100 (1981). 10.1007/BF00656997

48 Picard, M., White, K. & Turnbull, D. M. Mitochondrial morphology, topology, and membrane interactions in skeletal muscle: a quantitative three-dimensional electron microscopy study. J Appl Physiol (1985) 114, 161–171 (2013). 10.1152/japplphysiol.01096.2012

49 Linscheid, N. et al. Quantitative proteomics and single-nucleus transcriptomics of the sinus node elucidates the foundation of cardiac pacemaking. Nature Communications 10, 2889 (2019). 10.1038/s41467-019-10709-9

50 Tran, H. T. N. et al. A benchmark of batch-effect correction methods for single-cell RNA sequencing data. Genome Biol 21, 12 (2020). 10.1186/s13059-019-1850-9

51 Hafemeister, C. & Satija, R. Normalization and variance stabilization of single-cell RNA-seq data using regularized negative binomial regression. Genome Biol 20, 296 (2019). 10.1186/s13059-019-1874-1

52 Xie, C. et al. KOBAS 2.0: a web server for annotation and identification of enriched pathways and diseases. Nucleic Acids Res 39, W316–322 (2011). 10.1093/nar/gkr483

53 Szklarczyk, D. et al. The STRING database in 2023: protein-protein association networks and functional enrichment analyses for any sequenced genome of interest. Nucleic Acids Res 51, D638–d646 (2023). 10.1093/nar/gkac1000

54 Ghazizadeh, Z. et al. A dual SHOX2:GFP; MYH6:mCherry knockin hESC reporter line for derivation of human SAN-like cells. iScience 25, 104153 (2022). 10.1016/j.isci.2022.104153

